# Programmed cell death and stomatal density regulate anther opening in response to ambient humidity

**DOI:** 10.1101/2024.09.09.612018

**Authors:** Anna Kampová, Moritz K. Nowack, Matyáš Fendrych, Stanislav Vosolsobě

## Abstract

Anther dehiscence is the process that facilitates pollen release from mature anthers in flowering plants. Despite its crucial importance to reproduction, the underlying molecular regulations and the integration of environmental information remain poorly understood. Using controlled humidity treatments of *Arabidopsis thaliana* flowers, we show here that high humidity prevents anthers from opening. Lower and higher stomatal densities correlate with slower and faster dehiscence dynamics, respectively, suggesting that controlled transpiration regulates anther opening. Furthermore, analyses of subcellular markers revealed that specific anther tissues undergo a spatially controlled programmed cell death (PCD) process as anthers open. Notably, genetic inhibition of PCD networks delays, whereas precocious PCD induction promotes, anther dehiscence. Our data shed new light on the interplay between ambient humidity, controlled transpiration, and PCD processes in regulating timely pollen release in the model plant Arabidopsis.

## Introduction

Anther dehiscence, the process of anther opening leading to pollen release, is crucial for pollination and thus reproduction in seed plants. Due to its importance, anther dehiscence has been investigated for many years^1–3^. Already almost two centuries ago, Purkyně in his pioneering works described various anther tissues that are crucial for dehiscence in several species^4^. Since then, a wide range of plant species has been used for research, including many non-model species^5–7^. In general, different types of tissues can be distinguished in angiosperm anthers: tapetum, middle layer, endothecium, epidermis, septum and stomium^8,9^. Each tissue is essential and contributes to pollen and anther development or dehiscence. There are well-described developmental processes that precede dehiscence, such as tapetum degradation^10^ or secondary cell wall thickening in the endothecium^11^.

It has been proposed that dehiscence requires tissue dehydration^5,12^. Dehydration is facilitated by water transport within the anther^13,14^ and controlled transpiration via stomata^5^. As transpiration rates depend on external conditions, the effect of high humidity has been investigated, assuming that it interferes with anther opening. However, these experiments on various species like rice^15^, apricot or peach^16^, and pecan^17^, were performed on detached stamens or flowers, potentially creating artefacts. Although the impact of weather, particularly humidity, on dehiscence may offer plants a competitive advantage by selectively exposing pollen to pollinators^18,19^, the underlying mechanisms are not yet fully understood.

The water balance in plant tissues is generally regulated by alternation of stomata aperture, and anther dehydration is no exception. This is evident in the *Arabidopsis thaliana inducer of cfb expression1-2* (*ice1-2*) mutant, which failed to open anthers due to disrupted stomatal development^20^. In the context of transpiration, water transport is no less important. For example, tobacco (*Nicotiana tabacum)* plants with RNA-silenced aquaporin PIP2 exhibited delayed anther dehiscence, highlighting the relevance of controlled water movement^14^. In *A. thaliana* anthers, water transport is regulated by jasmonic acid (JA)^21^, as shown in mutants with disrupted JA biosynthesis *defective in anther dehiscence1* (*dad1*) and *delayed dehiscence1* (*dde1*) that display abnormal anther dehiscence due to lack of dehydration^22,23^.

Developmentally controlled programmed cell death (dPCD) of different tissues throughout anther development has been suggested as an important pathway in anther development^24,25^. dPCD in the tapetum has been reported as an essential process for both pollen and anther development in rice (*Oryza sativa*)^26,27^ and Arabidopsis^28^. In *N. tabacum*, genetic ablation of stomium and surrounding tissues inhibited dehiscence, implicating the necessity of timely and controlled stomium degeneration for anther dehiscence^29^. Nonetheless, the coordination of cell death with anther dehydration remains unclear.

This study presents a comprehensive analysis of anther dehiscence in *A. thaliana*. Our controlled laboratory experiments provide evidence that ambient humidity and transpiration control anther opening. Genetic and cell biology studies revealed that dehiscence is promoted by a defined developmental program involving spatially and temporally controlled progression of dPCD in specific anther tissues.

## Results

### Anther dehiscence is postponed by high humidity

We investigated the hypothesis that ambient humidity affects dehiscence, which provides a potential link between weather conditions and pollen release. We established *A. thaliana* as a model system to examine the impact of high ambient humidity (HH) on anther dehiscence (Fig. 1A-B), developing an experimental setup to control humidity under lab conditions (Fig. 1C). The primary advantage of this setup is the selective exposure to HH (100% relative humidity), exposing only inflorescence stems to HH while the rest of the plant remains at regular ambient humidity (AH, approx. 60%). Targeting specifically the inflorescences limits systemic effects and eliminates the need for detaching floral parts, leaving the examined plants intact.

**Figure 1.**
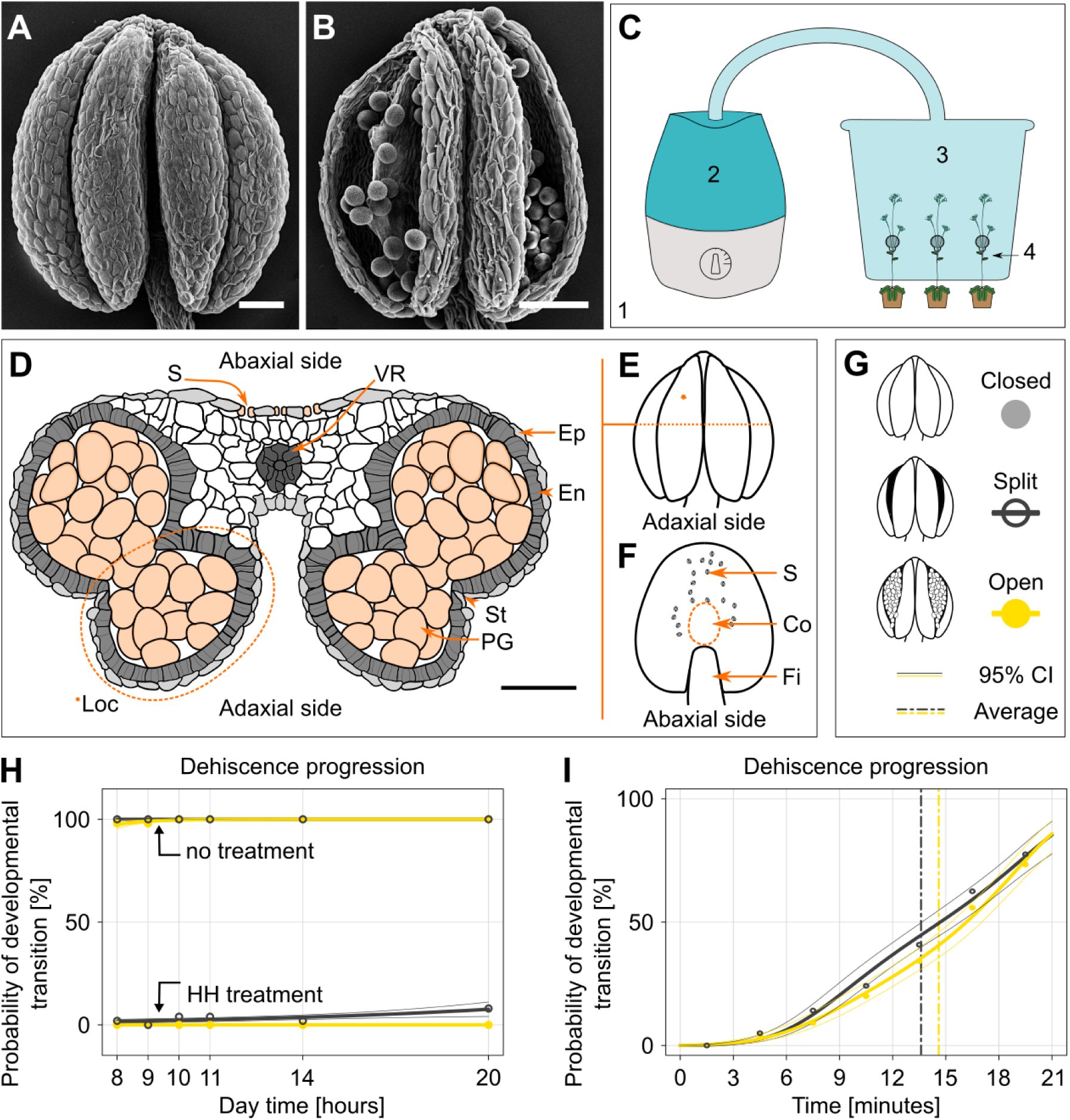
A. thaliana anthers do not open when exposed to high humidity (HH) in controlled conditions. **A.** Closed *A. thaliana* anther. **B.** Open *A. thaliana* anther. **C.** Cultivation chamber (*1*) with HH generator (*2*) creating HH conditions in a box (*3*) with openings for stem placement (*4*). **D.** Cross-section of a mature anther, *\*Loc*. anther locule. *S.* stoma, *VR.* vascular region, *Ep.* epidermis, *En.* endothecium with secondary thickenings, *St.* stomium, *PG.* pollen grains. The scheme is based on the confocal z-stack of *A. thaliana* anther cross-section obtained with a vibratome. **E.** Anther from the adaxial side; * indicates one of the four locules, and the dashed line indicates the cross-section. **F.** Abaxial side of the anther; *S.* wild-type stomata density and distribution, *Co.* connective, *Fi.* filament. **G.** Legend for the graphs K-L. Light grey represents a closed anther at stage 12 according to Sanders *et al*., 1999^8^. Dark grey stands for anther with a narrow slit formed at the apex – stage 13. Yellow is for fully open anther at stage 14a. Coloured points show measured proportions of respective developmental stages, logistic regression (GLM) is indicated by thick lines with 95% CI – thin lines. **H.** HH significantly prevents *A. thaliana* anthers from opening (*P* < 2.2e-16***, GLM logistic regression). 200 anthers were exposed to HH treatment from 5:00 to 20:00. Developmental stages were examined at 8:00, 9:00, 10:00, 11:00, 14:00 and 20:00. As a control in standard humidity, 200 anthers were analysed at the same time. Legend in G. **I.** Anther dehiscence was recorded over 21 minutes in 3-minute steps after the HH treatment finished at 11:00 (120 anthers). The dashed lines indicate the average values. Figures A and B were obtained using SEM. The scale bar is 50 µm in all figures.

Experiments always included only fully mature stamens of similar length as the pistil. The locules of mature anthers are surrounded by two cell layers, the epidermis and endothecium (Fig. 1D). The anthers of *A. thaliana* release pollen through two openings on the adaxial side, which are formed at the time of anther dehiscence (Fig. 1E). Of note, this adaxial side lacks stomata, while the abaxial side does form stomata (Fig. 1F). We designated three distinct categories to characterise anther dehiscence stages: closed (no visible opening), split (opening visible but not gaping at the anther apex), and fully open (pollen being released through gaping stomium) (stages 12, 13 and 14a, respectively, according to Sanders *et al.*, 1999^8^, Fig. 1G). *A. thaliana* flowers were exposed to HH from 5:00 in the morning onwards and dehiscence development was measured at 8:00, 9:00, 10:00, 11:00, 14:00 and 20:00. The HH-exposed anthers remained closed throughout the HH treatment (Fig. 1H, Tab. S1). Although at 20:00, 16 out of 200 anthers had split, none reached the fully open stage. In contrast, 196/200 anthers exposed to AH were fully open already by 8:00. The remaining 4 anthers were split at 8:00 and fully opened one hour later.

Next, we moved inflorescences to AH after 6 hours of HH treatment and measured anther dehiscence progression in 3-minute intervals for 21 minutes. Most of the analysed 120 anthers split and fully opened between 14 and 15 minutes after exposure to AH (Fig. 1I).

As stamen/flower detachment has been widely used in previous studies, we also analysed the effect of the detachment on dehiscence using the same experimental settings as described above. Contrary to intuition, detached flowers showed significant delays in anther dehiscence and full opening compared to attached flowers (Fig. S1A-D). This suggests that some dehiscence regulatory mechanisms, possibly involving stomatal function, require intact plants.

These experiments show that HH prevents anthers from splitting and full opening for a prolonged time and that dehiscence occurs within minutes after humidity decreases.

### Stomatal density affects anther dehiscence dynamics

As stomata control the gas exchange rate between the anther inner air spaces and the atmosphere and their dynamic aperture crucially determine transpiration rates, we tested whether stomatal density influences anther dehiscence, as had been previously suggested^20^. We analysed established *A. thaliana* mutant lines with higher stomatal densities (*epidermal patterning factor1,2* (*epf1,2*); *stomatal density and distribution1-1* (*sdd1-1*); and *too many mouths1-1* (*tmm1-1*), as well as a line with lower stomatal density (*STOMAGEN-EPFL9* (*stRNAi*)^30,31^. We confirmed that anthers in *epf1,2*; *sdd1-1*; and *tmm1-1* exhibited higher stomatal numbers and altered spacing compared to the WT, whereas *stRNAi* showed lower stomatal numbers (Fig. 2A-B). Next, we exposed inflorescences to HH to prevent dehiscence and then moved them to AH conditions. The progression of anther dehiscence was measured over 21 minutes and scored as described above. Anthers with increased stomatal numbers, specifically *epf1,2*; *sdd1-1*; and *tmm1-1*, demonstrated significantly faster full opening compared to the WT, whereas *stRNAi* anther opening occurred slower (Fig. 2C-H). Probabilities of transition from closed to split anthers were significantly higher for *epf1,2*; *sdd1-1*; and *tmm1-1* lines (43±3 %, 61±3 %, and 65±2 %, respectively) compared to the WT (29±2 %) and even splitting probability of *stRNAi* (10±2 %). Similarly, probabilities of full anther opening were significantly higher for mutant lines *epf1,2*; *sdd1-1*; and *tmm1-1* (38±3 %, 54±3 %, and 56±3 %) compared to WT (21±2 %) and much lower in *stRNAi* (6±1 %) (Fig. S2, Tab. S2).

**Figure 2:**
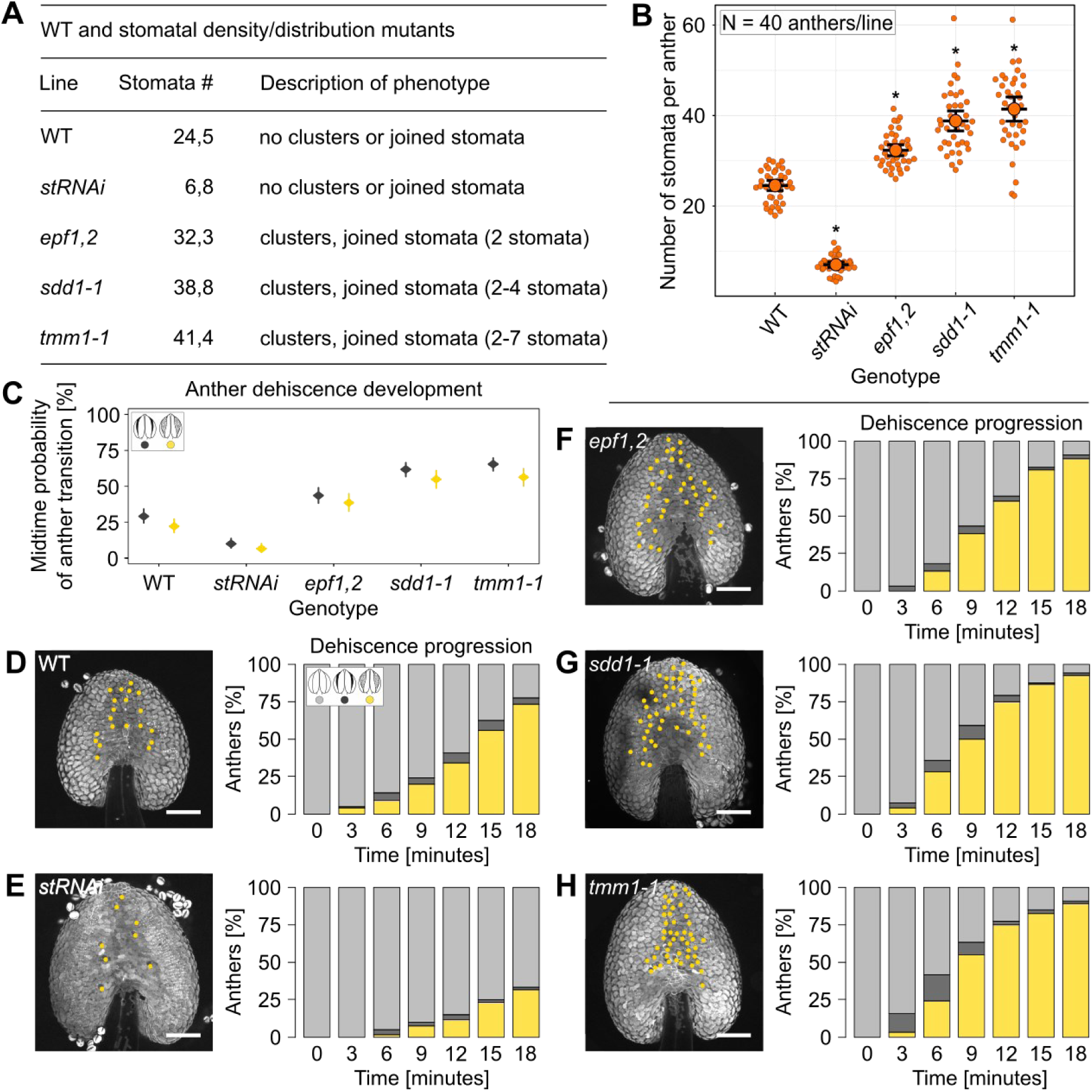
Anthers with higher stomata number and changed distribution (*epf1,2; sdd1-1; tmm1*) show accelerated opening compared to both wild-type (WT) and anthers with reduced stomata number (*stRNAi*). **A.** Table describing stomatal mutants. **B.** Number of stomata per average WT anther area. Median ± bootstrap 95% CI shown. Significant difference to WT is indicated by asterisk. **C-H. After high humidity treatment, anther dehiscence was measured over 21 minutes in 3-minute intervals.** In total, 120 anthers/line were analysed. **C.** Probability of anther splitting and opening in the midtime (10.5 minutes). In *stRNAi*, both anther splitting and opening rates are significantly reduced. In *epf1,2; sdd1-1; tmm1* splitting and opening rates increased (GLM, *P* < 2.2e-16***, rates and their 95% confidence intervals are presented). Legend in the figure. **D.** WT anther: At 21 minutes, 27 anthers closed, 5 partially open and 88 fully open. Legend in the figure applies on D-H. E. *stRNAi* anther: At 21 minutes, 80 anthers closed, 2 partially open and 38 fully open. F. *epf1,2* anther: At 21 minutes, 11 anthers closed, 3 partially open and 106 fully open. G. *sdd1-1* anther: At 21 minutes, 7 anthers closed, 2 partially open and 111 fully open. H. *tmm1-1* anther: At 21 minutes, 11 anthers closed, 2 partially open and 107 fully open. **Legend:** In the figures C and D-H: Light grey – closed anther – stage 12, dark grey – partially opened anther (splitting) – stage 13, yellow – fully opened anther stage 14a according to Sanders *et al.*, 1999^8^. Abaxial sides of anthers are presented, each yellow dot represents a stoma. Anther visualisation employed the autofluorescence of the cuticle using a blue channel on a spinning disc microscope, Zeiss Axio Observer.7/Yokogawa CSU-W1-T2. The scale bar is 100 µm in all figures.

In conclusion, stomatal number significantly influences anther dehiscence duration, with higher stomatal densities resulting in faster opening and less stomata leading to slower opening.

### Endodermis and endothecium remain fully viable until dehiscence begins

PCD has been implicated in anther development^24–26^. Therefore, we assessed the viability of the different tissues in the dehiscing anthers using available *A. thaliana* marker lines for different subcellular compartments: *pHTR5::NLS-GFP-GUS* for nuclei^32^, *p35S::PIP2-GFP* for the plasma membrane^33^, *pUBQ10::VAMP711-YFP* for the tonoplast^34^, and *pUBQ10::ToIM* for the integrity of the central vacuole^35^. Inflorescences were exposed to HH for 4 hours, stamens removed, and anthers imaged with a confocal microscope immediately (time 0) or 20 minutes after transfer from HH to AH. Both abaxial and adaxial sides were observed with the focus on epidermis and endothecium. Anthers imaged at time 0 were fully closed and epidermis and endothecium cells were viable, indicated by intact cellular compartments (Fig. 3A-F, S3-5A-F). 20 minutes after the exposure to AH, anthers were fully open and showed signs of cell death in both epidermis and endothecium (Fig. 3G-L), including vacuolar collapse (Fig. 3G-L, S5-6G-L), nuclear envelope breakdown (Fig. S3G-L), and plasma membrane endodomain shedding^36^ (Fig. S4G-L). Anther dehiscence time-lapses were also obtained confirming the above observations (Fig. S7).

**Figure 3:**
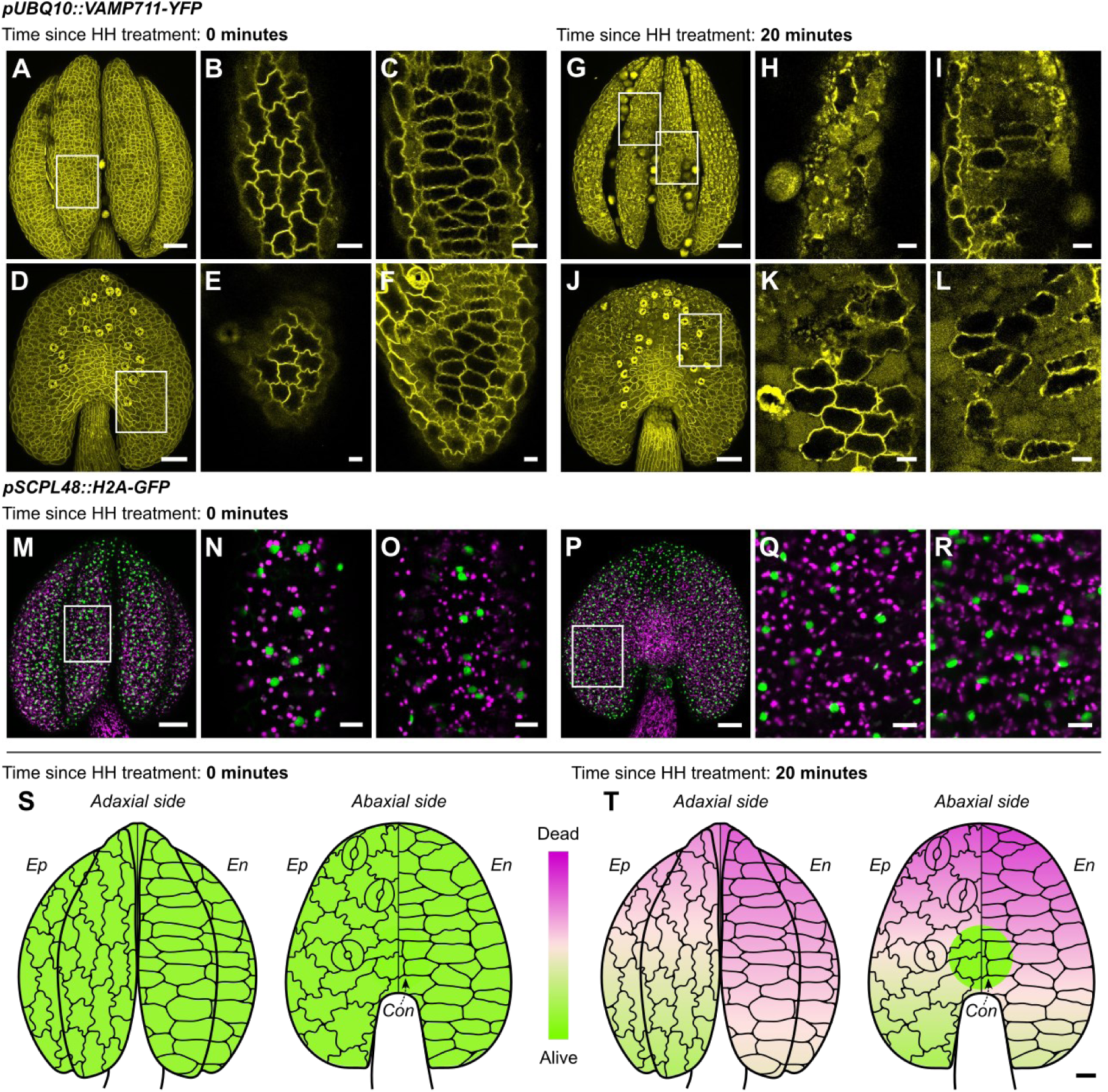
Anthers tissue – epidermis, endothecium and connective – are fully viable just before the opening starts. As the anther opens, the cells of mentioned anthers’ tissue undergo programmed cell death (PCD) in a direction-oriented manner. A-L: *A. thaliana pUBQ10::VAMP711-YFP* anthers are closed until the opening starts when short (4 hours) high humidity (HH) treatment is finished. Tonoplast degradation is visible 20 minutes into anther opening (tonoplast in yellow). Adaxial (AD, **A**) and abaxial side (AB, **D**) of closed anthers after short HH treatment, tonoplasts in both epidermis (**B, E**) and endothecium (**C, F**) are intact. AD (**G**) and AB side (**J**) of open anthers 20 min after short HH treatment, tonoplasts of the epidermis (**H, K**) and endothecium (**I, L**) are already degraded, but some cells remain intact. M-R: *pSCPL48::H2A-GFP* anthers show cells that are predetermined to undergo PCD. GFP, shown in green, autofluorescence in magenta. The expression includes the whole anther tissue: AD (**M**) and AB (**P**) sides, epidermis (**N, Q**) and endothecium (**O**, **R**). **S-T: PCD progresses from the anther apex to the base. Epidermal cell’s lifespan is longer than the endothecial cells. Connective tissue stays viable for the last.** In total, 112 anthers of these fluorescent marker lines were analysed: *pHTR5::NLS-GFP-GUS*, *p35S::PIP2-GFP, pUBQ10::ToIM*, *pUBQ10::VAMP711-YFP*, *pCEP1::H2A-GFP*, *pDMP4::H2A-GFP* and *pSCPL48::H2A-GFP*. Anthers were exposed to short HH treatment and observed 0 or 20 minutes after the HH treatment finished. Magenta represents completely dead cells, lawn green fully living cells and mistyrose represents the transition between dead and living tissue. Each anther scheme has an epidermis (*Ep*) on the left side and an endothecium (*En*) on the right side. The cell schemes represent the assigned tissue of respected anther sides, AD and AB. *Con.* indicates connective tissue. The cells are approximately 4× bigger in comparison to anther size. The data were analysed using beta regression in R (package **betareg**) **M.** Fully intact short HH treated anthers at the time of finished treatment. Epidermis and endothecium are fully viable on both AD and AB sides. **N.** Short-treated anthers 20 minutes after the treatment. PCD proceeds faster in the endothecium on both AD and AB sides. Anther apex contains more dead cells than the base. Connective is still viable. **Legend:** HH treatment – 4 hours. The figures were obtained using Leica TCS SP8. Scale bars are 50 µm in the anther overall view figures and 10 µm in close-up figures displaying cells. The scale bar for the anthers scheme is 10 µm and applies to the cells.

In summary, cells in the anther epidermis and endothecium are fully viable just before opening, but a rapid cell death occurs in these tissues as anther opening progresses.

### Cell death in anthers is a form of dPCD and progresses in a reproducible, directional pattern

To characterise the cell death processes during dehiscence further, we examined promoter-reporter lines of dPCD-upregulated signature genes (*pCEP1::H2A-GFP*, *pDMP4::H2A-GFP*, and *pSCPL48::H2A-GFP*)^37^. HH-treated plants were inspected as described above. At time 0 right after HH treatment, expression of nuclear-localized GFP of all dPCD-reporters was detected in the whole anther tissue, including the upper filament (Fig. 3M-R, S8-9A-F) suggesting a transcriptional preparation of dPCD in these tissues. Furthermore, signal diffusion indicative of chromatin degradation^38^ was observed after 20 minutes in AH (Fig. S8-10G-L). These results show that the rapid cell death observed in the epidermis and the endothecium during dehiscence has transcriptional parallels to established dPCD processes analysed in other tissues, suggesting that this cell death process is a form of dPCD. This conclusion is supported by the analysis of several dPCD signature genes in publicly available RNA-seq data (*CEP1*, *DMP4*, *SCPL48* mentioned above and *BFN1*, *EXI1*, *PASPA3*, *RNS3*). Our analysis shows that the expression of all these genes, except *PASPA3*, is upregulated during final stages of anther development, further indicating that a canonical dPCD occurs during anther dehiscence (Fig. S11A-G, Tab. S3-4).

To analyse the precise spatial pattern of dPCD progression during dehiscence, we analysed confocal z-stack images of several subcellular marker lines (*pHTR5::NLS-GFP-GUS*, *p35S::PIP2-GFP, pUBQ10::VAMP711-YFP*, *pUBQ10::ToIM*, *pCEP1/pDMP4/pSCPL48::H2A-GFP*; 8 anthers imaged per line after 4 hours of HH treatment, and 20 min after shift to AH). Each anther area was divided into regions and scored based on viability status, ranging from completely intact cells to decompartmentalised cells (Fig. S12I-J) in the epidermis and the endothecium. Data were then fitted using beta regression (Tab. S5). Our analyses revealed distinct differences between closed and fully open anther tissues, with closed anthers exhibiting a complete viability (Fig. 3S) and fully open ones containing dead cells in both epidermis and endothecium (Fig. 3T). By contrast, cells close to the connective area retained viability even after full anther opening. In both the epidermis and the endothecium, dPCD progression followed a basipetal pattern, with the anther base remaining viable the longest. Interestingly, we observed differences between the epidermis and endothecium: while the dPCD progression occurred in the same directional pattern in both tissues, dPCD in the endothecium spread faster toward the anther base than dPCD in the epidermis (Fig. 3T).

Taken together, the expression of dPCD signature genes, the rapid cellular decompartmentalization, and the distinctive pattern of cell death events suggest that a developmentally controlled, environmentally triggered dPCD process occurs during anther dehiscence.

### dPCD manipulation modulates anther dehiscence

Building on the dPCD observations during anther dehiscence, we hypothesized that dPCD promotes anther opening. We analysed the link between dPCD and anther dehiscence by exposing the above-mentioned marker lines (*pHTR5::NLS-GFP-GUS*, *p35S::PIP2-GFP, pUBQ10::VAMP711-YFP*, *pUBQ10::ToIM*, *pCEP1/pDMP4/pSCPL48::H2A-GFP*) to 28 hours of HH (long treatment) before transferring them to AH, artificially delaying dehiscence by one day. Confocal imaging immediately after this long treatment revealed that the endothecium contained cells that had undergone dPCD, while the entire epidermis layer remained viable (Fig. 4A, S3-6M-Y, 8-10M-Y, S12F). After 20 minutes in AH, dPCD occurred in both cell layers, although more rapidly in the endothecium (Fig. S12G-H). Dehiscence in WT anthers exposed to long HH treatment showed a splitting and full-opening probability of 48±3% and 56±3%, respectively, 10.5 minutes after the transfer from HH to AH. This probability is twice that of short-HH-treated (4 hours) WT anthers after the transfer from HH to AH (Fig. 4B, S13A-B, Tab. S6), showing that extended HH treatment delays, but does not altogether block, dPCD in anther tissues, correlated with anther dehiscence dynamics under these conditions.

**Figure 4:**
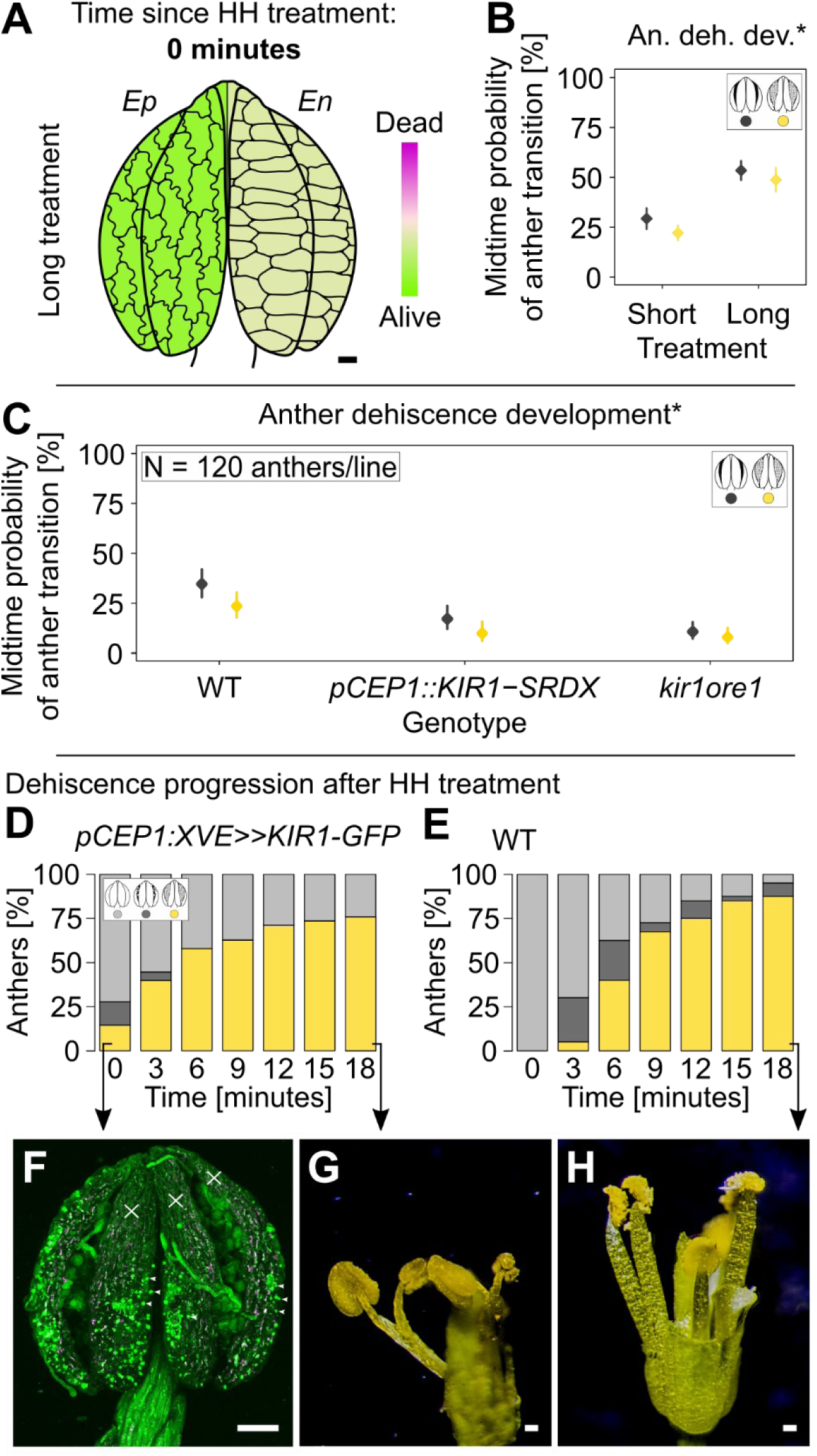
The programmed cell death (PCD) timing can be manipulated, resulting in changed anther dehiscence progression. Delayed PCD leads to slower opening. Earlier induced PCD execution results in opening despite high humidity treatment. **A-B.** Anthers were exposed to long HH treatment (28 hours). **A.** Long-HH-treated anthers of listed lines were imaged and analysed at the time of finished treatment: *pHTR5::NLS-GFP-GUS*, *p35S::PIP2-GFP, pUBQ10::ToIM*, *pUBQ10::VAMP711-YFP* and *pCEP1/pDMP4/pSCPL48::H2A-GFP*. The epidermis is still intact, but the endothecium undergoes PCD. Lawn green shows fully living tissue, and mistyrose represents tissue where PCD occurs but is not completely dead yet. The data were analysed using beta regression in R (package **betareg**) **B.** Dehiscence was measured over 21 minutes in WT short (4 hours) and long HH-treated anthers. Midtime opening and splitting probability is higher in WT long-HH-treated anthers (GLM, *P* < 2.2e-16***, rates and their 95% confidence intervals are presented). **C.** Anther dehiscence was measured over 21 minutes in lines with reported delayed PCD, *pCEP1::KIR1-SRDX* and *kir1ore1*. Midtime opening and splitting probability is lower in both lines than in WT (GLM, *P* < 2.2e-16***, rates and their 95% confidence intervals are presented, a legend in the figure). **D-H.** Anther dehiscence was measured over 21 minutes in *pCEP1::XVE≫KIR1-GFP* and WT, treated with β-estradiol 24 hours before the measurement. **D.** Bar plots show the ratio of closed, split and open anthers. At the time 0, 23 out of 83 *pCEP1::XVE≫KIR1-GFP* anthers are already split or open despite short high humidity (HH) treatment. Legend in the figure. **E.** All 40 WT anthers are fully closed at the time 0. Legend in D. **F.** At the time 0, *pCEP1::XVE≫KIR1-GFP* anther is open despite the HH treatment. Areas of dead tissue are marked with crosses and nuclei are indicated by asterisks. G. *pCEP1::XVE≫KIR1-GFP* flower after 21 minutes, anthers are open and filaments are shrunk. **H.** WT flower after 21 minutes, anthers are open and filaments are fully turgid. **Legend:** Figure F was obtained with Leica TCS SP8, the scale bar here is 50 µm. Figures G-H are from a stereomicroscope STM 822 5410 with Nikon D3200 camera, scale bars are 100 µm. In Figure A, the scale bar is 10 µm and applies to the cells that are approximately 4× times bigger than anther size.

To further test our hypothesis, we employed different approaches to either delay or induce dPCD by manipulating two established dPCD-promoting NAC transcription factors, *KIRA1* (*KIR1*) and *ORESARA1* (*ORE1*)^39^, which we selected as we found them to be expressed in later stages of anther development, based on publicly available expression data (Fig. S11H-I).

First, we measured dehiscence after exposure to HH followed by transfer to AH in mutant lines that have been shown to also affect stigmatic papilla lifespan^39^. Both the *kir1ore1* double mutant as well as dominant-negative *pCEP1::KIR1-SRDX* lines exhibited a significant delay in dehiscence. At 10.5 minutes after removal from HH chamber, the WT full opening probability was 27±3 %, compared to 8±2 % in *kir1ore1* and 10±2 % in *pCEP1::KIR1-SRDX* (Fig. 4C, S14A-D, Tab. S6). This identified a role of NAC-dependent dPCD gene regulatory networks during anther dehiscence and suggested that dPCD progression is required for fast anther opening after exposure to AH.

In a complementary approach, we induced precocious dPCD using an established estradiol-inducible overexpression of KIR in *pCEP1::XVE≫KIR1-GFP* line (Fig. S15)^39^. First, by analysing *pCEP1::H2A-GFP* anthers, we confirmed the necessary activity of *CEP1* promotor one day before anthesis (Fig. S14E-J). Next, the *pCEP1::XVE≫KIR1-GFP* anthers were treated with an estradiol solution at stage 11 exactly 24 hours before the anticipated dehiscence (Fig. S14K). Anthers were then exposed to HH before transfer to AH as described above. Whereas the overall progress of dehiscence after the transfer to AH did not significantly differ between the induced and control variants (Fig. S14S-T, Tab. S7), a significant proportion of induced anthers fully opened even under HH conditions, forming slits from apex to base, while control anthers remained closed in HH (Fig. 4D-E). Induced *pCEP1::XVE≫KIR1-GFP* anthers, that fully opened despite HH, contained partially dead tissue, especially at the anther apex (Fig. 4F, S14O-R). Moreover, induced pCEP*1::XVE≫KIR1-GFP* filaments were visually shrivelled (Fig. 4G), while control filaments appeared macroscopically turgid and viable at anther dehiscence (Fig. 4H), confirming the efficiency of the treatment to induce dPCD. These observations show that dPCD induction can overcome the inhibitory effect of HH on dehiscence and lead to anther opening even under HH conditions. However, dPCD alone seems not sufficient to enable the full opening and release of pollen grains, which necessitates AH to promote effective tissue dehydration.

In conclusion, by employing three different approaches, we demonstrate that dPCD processes in the anther epidermis and endothecium tightly correlate with anther dehiscence and that dPCD manipulation modulates dehiscence dynamics, strongly suggesting that dPCD processes in the anther epidermis and endothecium are a central part of an effective anther dehiscence mechanism.

## Discussion

### Environmental humidity regulates dehiscence and pollen release

Anther dehiscence is a process that occurs at the end of anther development and is crucial for timely and effective pollen release. It has been reasoned that it would be beneficial to reproductive success if pollen would be displayed to pollinators when they are most active, and for many plants that would be during the day and under dry weather conditions^18^. Dehiscence results from dehydration of anther tissue, presumably facilitated by aquaporin-promoted water transport within the anther tissue^14^ and controlled transpiration through stomata^5,25^.

These observations propose that anther opening is not a passive dehydration process, but closely controlled by the plant in response to ambient conditions. Therefore, we investigated the effect of this weather parameter, mimicked by high humidity in strictly controlled laboratory conditions on intact plants and flowers. We show that high ambient humidity strongly affects anther opening, postponing both splitting and full opening. A drop in humidity triggers rapid anther dehiscence as the anthers fully open within 15 minutes after exposure to lowered ambient humidity. During this time, dehydration of the epidermis changes its mechanical properties resulting in complete anthers’ shape change^40^.

Our conclusions are supported by a diversity of earlier observations from plant species such as rice^15^, apricot, peach, almond^16^, and pecan^17^. These were, however, conducted on dissected flowers or stamens and we observed that dissection of flowers led to delayed anther dehiscence compared to dehiscence in anthers of intact plants. As a drought stress response may be triggered in the detached plant organs, it is tempting to speculate that the drought-induced plant hormone abscisic acid may negatively influence anther opening by causing stomata to close. Importantly, our results on Arabidopsis flowers reveal that anther opening in response to lowered ambient humidity can a very rapid process, which necessitates analyses with high temporal resolution to dissect the mechanisms involved.

Our observations strongly support the notion that plants can actively control dehiscence, as also suggested by dynamic changes in phytohormone levels observed in *Solanum lycopersicum*^41^. To verify the effect of humidity and transpiration on dehiscence genetically, we examined various stomata mutants. Mutants with higher stomata number, *epf1,2*; *sdd1-1*; and *tmm1-1,* showed faster dehiscence, whereas *stRNAi* anthers with the least number of stomata displayed the most delayed dehiscence. This is in line with the earlier observations that *A. thaliana ice1-2* anthers exhibit disrupted dehiscence correlating with defective stomata development^20^.

In summary, our controlled selective humidity treatments of intact flowers revealed that dehiscence is strongly delayed in high ambient humidity conditions and rapidly triggered by humidity decrease. Requirement of functional stomata further confirms that transpiration is a necessary part of a controlled anther dehiscence mechanism. These findings make it tempting to speculate that the stomata formed on the abaxial side of anthers are primarily important to facilitate a rapid tissue dehydration leading to anther opening.

### Dehiscence is developmental program depending on dPCD

Previous studies have shown the involvement of dPCD in specific aspects of anther development, for instance tapetum maturation^27^. Also the degeneration of stomium cells during anther development has been observed and implicated in anther dehiscence as experiments suggest that dPCD processes in this tissue lead to gaping stomium^42^. Our investigations revealed that in addition to controlled transpiration and water transport, also a rapidly triggered dPCD process in epidermis and endothecium plays a role in effective and rapid anther opening.

Analyses of various fluorescent marker lines revealed that the anthers stayed closed, and epidermis and endothecium cells remained viable under high humidity conditions, supporting observations in the anther tissue in lily^43^. After the transfer to ambient humidity, the anthers fully opened, while first endothecium and then epidermis cells underwent rapid cell death in a basipetal fashion starting at the anther apex. Dying cells displayed features of cellular decompartmentalization typical for dPCD, with nuclei and vacuoles collapsing, and the plasma membrane exhibiting endodomain shedding^36^. In addition, we detected the expression of dPCD signature genes^37^, as well as established transcription factors regulating dPCD gene regulatory networks^38,39^ in pre-dehiscence anthers. The subsequent evaluation of all marker lines showed a specific spatial pattern of dPCD progression in anthers.

We used genetic manipulation of dPCD to examine its importance for anther dehiscence using lines with delayed dPCD, *kir1ore1* and *pCEP1::KIR1-SRDX*^39^. Both lines exhibited delayed dehiscence compared to the WT. In a complementary approach, the induction of the precocious dPCD in the *pCEP1::XVE≫KIR1-GFP* revealed that dPCD is sufficient to promote dehiscence. Anthers with induced dPCD showed fully opened slit along the whole axis and partially exposed pollen grains even in high humidity. This shows that dPCD is required and sufficient for dehiscence; specifically, for the anther splitting.

As cells undergo dPCD, they shrink^44^ and even small changes in turgor can induce large movements, depending on tissue characteristics^45^. However, the overall shape of anthers after precocious dPCD under HH had been induced was still maintained, supporting the necessity of transpiration and evaporation in addition to dPCD to achieve efficient and rapid anther dehiscence. According to a model based on lily and Arabidopsis, dehydration of the epidermis, together with secondary endothecial thickening, are the driving mechanical forces that facilitate the movement of the anther tissues from closed to open^40^. Our analysis shows that dPCD in the epidermis and endothecium is sufficient for partial anther opening under HH conditions, and necessary for rapid anther opening under AH conditions. Therefore, dPCD may on one hand accelerate dehydration processes and on the other contribute to tissue deformation through rapid loss of turgor pressure after cell death.

To conclude, anther dehiscence is a tightly regulated process that integrates a developmentally prepared PCD program with a rapid response to ambient humidity, providing a mechanism to optimize pollen release. Interestingly, a rapid onset dPCD execution appears to be triggered by low ambient humidity. The underlying mechanism of the dPCD regulation by humidity, as well as control of stomata aperture to facilitate tissue dehydration are intriguing topics for future investigations.

## Materials and Methods

### Plant material and growth conditions

For this study, we used the following wild-type line: *A. thaliana* Col-0 ecotype, *A. thaliana* fluorescent marker lines used were: *p35S::PIP2-GFP*^33^, *pHTR5::NLS-GFP-GUS*^32^, *pUBQ10::ToIM*^35^ and *pUBQ10::VAMP711-YFP*^34^. PCD-promoter marker lines used were *pCEP1::H2A-GFP*, *pDMP4::H2A-GFP* and *pSCPL48::H2A-GFP*^37^. Other PCD-related lines were employed: *kir1ore1*, *pCEP1::KIR1-SRDX* and *pCEP1::XVE≫KIR1-GFP*^39^. Stomata mutants used were *epf1,2*; *tmm1-1*; *stRNAi*^30^ and *sdd1-1*^31^. All the lines are listed in Tab. S8. The seeds were sown on peat pellets (Jiffy). Fertilizer Kristalon Gold was used for the first watering (1 g of fertilizer per 1 L of water). Pellets with seeds were placed at dark and 4 °C for 7 days. Plants were then grown in long-day conditions (16 hours light and 23 °C / 8 hours darkness and 18 °C; light intensity was 120 μmol m^-2^ ^s-1^).

### High humidity treatment and anther dehiscence measurement

High humidity conditions were generated by the Lucky Reptile SuperFog II dew generator. This machine was connected through a plastic hose with a transparent plastic box (38×28×28 cm, Ikea). The box was closed by a lid which had an opening in it to put the hose which led the wet air from the machine to the box (Fig. 1C). Number of bigger and smaller holes were made in all the box sides at the height of grown *A. thaliana* plants. Through the bigger hole, a stem with the closed buds was placed inside and then moved to the smaller one down below. The bigger hole was then secured with a foam plug to reduce the high humidity leakage. In the evening prior the experiment, old flowers were removed from the inflorescence, which was then placed inside the box. Treatment started at 5:00 and finished according to the experimental setup. To monitor the temperature and humidity in the HH box, an automatic system based on Raspberry Pi Zero W was developed using one Adafruit TSL2591 light sensor and two Adafruit BME280 temperature/humidity sensors placed in and out of the HH box. The relative air humidity inside the box during the treatment was 100 %. The fully saturated air led to water condensation on the plant surface. The ambient humidity outside the box was approximately 60 %. The stage of anther dehiscence was recorded once per hour and placed back into the box (Fig. 1H) or the stems were removed from the box and the anther dehiscence progress was measured for 21 minutes in 3-minute steps (Fig. 1I, 2C-H, 4B-C and D-E, S1B-D, S2, S13A-B, S14A-D and S-T) using a binocular stereomicroscope. The stages of anther opening were scored according to Sanders *et al*., 1999^8^: anthers closed – stage 12; anthers partially open – stage 13; anthers open – stage 14a (Fig 1G), the ambient temperature was measured by a mercury thermometer.

### Gene expression induction

The gene induction was based on Gao *et al.*, 2018^39^ protocol. For this experiment, flower buds one day before anthesis were chosen. At this developmental stage, anthers were already yellow and reached more than half of the pistil size (Fig. S13K). All the sepals and petals were removed from the flower together with a pistil. All unmatured buds were also removed from the stem. The stem was then snapped to a wooden skewer and the flower was placed in an upside-down PCR tube also snapped to the skewer. The tube was filled with 350 µL of a working solution containing 1/20 MS and 100µM β-estradiol in DMSO (β-estradiol DMSO-based stock was 20mM). The treatment lasted 6 hours and was done one day before anthesis exactly 24 hours before the anther opening measurement.

### Confocal and light microscopy

For confocal microscopy imaging, Stamens were attached to double-sided tape by their filaments and embedded in 2,5% low-gelling agarose (Sigma-Aldrich) and a covered slide was placed on top. All the figures (except for Fig. 1A-B and Fig. 2D-H) were obtained using a confocal microscope Leica TCS SP8 with the 20x/0.75 IMM objective using water immersion. The excitation and emission spectra were as follows: GFP λ_EX_ = 488 nm, λ_EM_ = 495-540 nm or 495-550 nm; chloroplasts autofluorescence 561 nm, λ_EM_ = 614-749 nm; ToIM - GFP λ_EX_ = 488, λ_EM_ = 495-540 nm, tagRFP λ_EX_ = 561 nm, λ_EM_ = 570-660 nm; YFP λ_EX_ = 514 nm, λ_EM_ = 520-580 nm.

The figures for Fig. 2 were obtained using a vertical stage^46^ spinning disc microscope Zeiss Axio Observer.7/Yokogawa CSU-W1-T2 with a VS-HOM1000 excitation light homogenizer and with the use of Plan-Apochromat 20x/0.8 M27 objective. The autofluorescence of the cuticle was visualized using a blue channel, λ_EX_ = 405 nm. For time-lapse microscopy, plants were transferred to the microscopy room at 5:40 in the morning, with the avoidance of their exposure to light. Under the stereomicroscope with the lowest possible illumination, the inflorescence, while still connected to the plant, was attached to the microscope slide by commercial double-side adhesive tape. The flower bud that was expected to open on a given morning was carefully opened by tweezer and both petals and sepals were also fixed to the tape. Eventually, the stigma and some of the pistils were removed for better observation. The slide together with the whole plant was placed in the vertical stage spinning disk microscope and observed as a dry mounting (λ_EX_ was 488 and 561 nm for the *pUBQ10::ToIM* line and 515 nm for *pUBQ10::VAMP711-YFP*, Plan-Apochromat 10×/0.45 M27, z-step 7-10 μm, time step 30-60 s). Brightfield figures were acquired using the Nikon D3200 camera and stereomicroscope STM 822 5410.

### Scanning electron microscopy

Phosphate buffer (PBS) of pH 7.2 was prepared for SEM sample preparation. First Solution A was made using 7.146 g Na_2_HPO_4_. 12 H_2_O for 100 mL and 2.76 g for 100 mL of NaH_2_PO_4_. H_2_O for solution B. Solutions were mixed to create 0.2M stock of PBS; 36 mL of solution A and 14 mL of solution B were used. Samples were fixed in a solution of 2.5% glutaraldehyde (Grade I for electron microscopy, Sigma-Aldrich) in 0.1M PBS for 24 h at 4 °C. After the fixation, samples were washed with 0,1M PBS for 12 h at 4 °C and then with distilled water for 10 min at RT. Dehydration by EtOH series with increasing concentration followed: samples were placed in EtOH of concentration 35% for 15 min, 50% for 15 min, 70% for 30 min, 80% for 15 min, 96% for 15 min and finally 100% for 15 min. Samples were transferred to paper envelopes while kept wet in 70% EtOH, they were then dried in a critical point dryer Bal-Tec CPD 030. Samples were coated by a 2 nm gold layer in an ion sputter coater Bal-Tec SCD 050 and observed with a JEOL JSM-IT200 HR scanning electron microscope.

### Quantification of cell death progression in anthers

Analysis of PCD progression was made on both short and long humidity-treated anthers for fluorescent marker lines *pHTR5::NLS-GFP-GUS*, *p35S::PIP2-GFP, pUBQ10::ToIM*, *pUBQ10::VAMP711-YFP*, *pCEP1::H2A-GFP*, *pDMP4::H2A-GFP* and *pSCPL48::H2A-GFP*. The plants were exposed to HH conditions as described above. The short treatment lasted for 4 hours from 5:00 to 9:00, the long treatment lasted for 28 hours from 5:00 day 1 to 9:00 day 2. Anthers were imaged by a confocal microscope immediately after the HH treatment or 20 minutes later with the dehiscence already progressing. The proportion of dead cells in the anther tissue was assessed semiquantitatively using categories of > 10 %, 10-40 %, 40-60 %, 60-90 % and > 90 % dead cells. The apical, central and basal regions (and connective, if applicable) were assessed separately for the adaxial and abaxial sides and both epidermis and endothecium (Fig. S12 I-J). In the apical, central and basal horizontal regions in the adaxial side, the four locules were scored separately and then averaged. In the abaxial side, central and basal horizontal regions were scored left and right separately and then again averaged. The reliability of scoring was enhanced by horizontal averaging of corresponding areas. The dependence of the spatial proportion of dead cells on dehiscence progression and length of previous humidity treatment was fitted by beta regression in R^47^ (package **betareg**), the optimal model was selected using the StepBeta function (package **StepBeta**), and the significance of specific variables was determined by likelihood ratio test (lrtest, package **lmtest**). The complete R script (CellDeathQuant.md) can be found in our GitHub repository. In the analysis, 112 anthers were included, and 370 anthers were imaged in total. The latter number also represents anthers from all the time points and lines which were not included.

### Statistical analysis of anther openings

Logistic regression of HH treatment experiments on *A. thaliana* was analysed by GLM with binomial distribution and individual effect significance was calculated by χ^2^-test. 95% confidence intervals were calculated by `confint` function with default setting.

Anther opening after HH treatment was analysed by logistic regression (GLM with binomial family of distribution). Variant identifiers, time and its quadrate were used as predictors in a model with a full set of interactions. Optimal model was determined by a stepwise algorithm according to AIC (`step` function) and tested by χ^2^-test (`anova` function). Differences between individual variants were estimated by `emmeans` function from package **emmeans**. PlotsOfData web app^48^ was used to plot the data on the stomata number (Fig. 2B).

### Analysis of PCD-related genes expression

The set of RNA-seq libraries from 12 different stages of *A. thaliana* anther, 3 transcriptomes of mature pollen and one transcriptome of filament was obtained from the ENA repository (in total 22, 10 and 2 libraries, Tab. S3) and quantified by Kallisto 0.48.0 using Araport11 representative CDS model as a reference (Tab. S4). For other libraries, non-standardized TPM values of transcripts of interest were fitted by the GAM model (R package **mgcv**^49^), while medians were counted in the case of pollen and filament libraries.

### Image analysis

The brightfield image series were focus-stacked in an app PICOLAY (www.picolay.de) and then adjusted with a program Zoner Photo Studio X (https://www.zoner.cz). All the figures were processed using Fiji^50^. For stomata number analysis, a newly developed Fiji macro surface_.ijm was utilized to generate the surface figures. For time-lapse imaging, acquired 16-bit z-stacks were processed also in Fiji subjected to maximal projection and converted to 8-bit AVI after nonlinear brightness adjustment (RawTherapee 5.8, PP3 conversion files are in supplement). Every figure was assembled in Inkscape^51^.

## Supporting information

Supplementary file PP3 conversion

## Acknowledgement

The authors are grateful to Marie Hronková (USB České Budějovice, Czechia) for providing the seeds of stomata mutant lines. The authors thank Petra Cifrová, Eliška Kobercová, Jan Martinek and Josef Šonka for advice and support.

## Author contributions

AK carried out the experiments and wrote the manuscript. SV analysed the data with AK input. MF offered valuable consultation throughout the project, bringing insights that significantly shaped the work’s direction. MN provided the seeds of the PCD marker lines, PCD inducible lines, and PCD delayed lines, and gave valuable feedback and comments on the manuscript.

## Data and code availability

Source codes for all analyses are available in the GitHub repository: https://github.com/vosolsob/anthers/.

## References

1. Keijzer, C. J. Mechanisms of Angiosperm anther dehiscence, a historical review. Anther and Pollen 55–67 (1999) doi:10.1007/978-3-642-59985-9_6.

2. Venkatesh, C. S. The form, structure and special ways of dehiscence of anthers of Cassia—III. Subgenus Senna. Phytomorphology 7, 253–273 (1957).

3. Zhao, S. Q., Li, W. C., Zhang, Y., Tidy, A. C. & Wilson, Z. A. Knockdown of Arabidopsis ROOT UVB SENSITIVE4 Disrupts Anther Dehiscence by Suppressing Secondary Thickening in the Endothecium. Plant Cell Physiol. 60, 2293–2306 (2019).

4. Purkyně, J. E. De cellulis antherarum fibrosis nec non de granorum pollinarium formis: commentatio phytotomica. (1830).

5. Keijzer, C. J. The processes of anther dehiscence and pollen dispersal: I. The opening mechanism of longitudially dehiscing anthers. New Phytol. 105, 487–498 (1987).

6. Wang, D. K., Sun, G. F., Wang, L. F., Zhai, S. & Cen, X. J. A novel mechanism controls anther opening and closing in Paris polyphylla var. Yunnanensis. Chinese Sci. Bull. 54, 244– 248 (2009).

7. Lyu, X. et al. Characterization of watermelon anther and its programmed cell death-associated events during dehiscence under cold stress. Plant Cell Rep. 38, 1551–1561 (2019).

8. Sanders, P. M. et al. Anther developmental defects in Arabidopsis thaliana male-sterile mutants. Sex. Plant Reprod. 11, 297–322 (1999).

9. Xue, J. S. et al. Development of the Middle Layer in the Anther of Arabidopsis. Front. Plant Sci. 12, 1–9 (2021).

10. Yi, J. et al. Defective Tapetum Cell Death 1 (DTC1) Regulates ROS Levels by Binding to Metallothionein during Tapetum Degeneration. Plant Physiol. 170, 1611–1623 (2016).

11. Dawson, J. et al. Characterization and genetic mapping of a mutation (ms35) which prevents anther dehiscence in Arabidopsis thaliana by affecting secondary wall thickening in the endothecium. New Phytol. 144, 213–222 (1999).

12. Bonner, L. J. & Dickinson, H. G. Anther dehiscence in Lycopersicon esculentum Mill. I. Structural aspects. New Phytol. 113, 97–115 (1989).

13. Stadler, R., Truernit, E., Gahrtz, M. & Sauer, N. The AtSUC1 sucrose carrier may represent the osmotic driving force for anther dehiscence and pollen tube growth in Arabidopsis. Plant J. 19, 269–278 (1999).

14. Bots, M. et al. Aquaporins of the PIP2 class are required for efficient anther dehiscence in tobacco. Plant Physiol. 137, 1049–1056 (2005).

15. Matsui, T., Omasa, K. & Horie, T. Mechanism of anther dehiscence in rice (Oryza sativa L.). Ann. Bot. 84, 501–506 (1999).

16. Gradziel, T. M. & Weinbaum, S. A. High relative humidity reduces anther dehiscence in apricot, peach, and almond. HortScience 34, 322–325 (1999).

17. Yates, I. E. & Sparks, D. Environmental regulation of anther dehiscence and pollen germination in pecan. J. Am. Soc. Hortic. Sci. 118, 699–706 (1993).

18. Štenc, J. et al. Pollinator visitation closely tracks diurnal patterns in pollen release. Am. J. Bot. 110, 1–11 (2023).

19. Bassani, M., Pacini, E. & Franchi, G. G. Humidity stress responses in pollen of anemophilous and entomophilous species. Grana 33, 146–150 (1994).

20. Wei, D. et al. INDUCER OF CBF EXPRESSION 1 is a male fertility regulator impacting anther dehydration in Arabidopsis. PLOS Genet. 14, e1007695 (2018).

21. Acosta, I. F. & Przybyl, M. Jasmonate Signaling during Arabidopsis Stamen Maturation. Plant Cell Physiol. 60, 2648–2659 (2019).

22. Sanders, P. M. et al. The Arabidopsis DELAYED DEHISCENCE1 gene encodes an enzyme in the jasmonic acid synthesis pathway. Plant Cell 12, 1041–1062 (2000).

23. Ishiguro, S., Kawai-Oda, A., Ueda, J., Nishida, I. & Okada, K. The DEFECTIVE IN ANTHER DEHISCENCE1 Gene Encodes a Novel Phospholipase A1 Catalyzing the Initial Step of Jasmonic Acid Biosynthesis, Which Synchronizes Pollen Maturation, Anther Dehiscence, and Flower Opening in Arabidopsis. Plant Cell 13, 2191–2209 (2001).

24. Koltunow, A. M., Truettner, J., Cox, K. H., Wallroth, M. & Goldberg, R. B. Different temporal and spatial gene expression patterns occur during anther development. Plant Cell 2, 1201–1224 (1990).

25. Wilson, Z. A., Song, J., Taylor, B. & Yang, C. The final split: The regulation of anther dehiscence. Journal of Experimental Botany vol. 62 1633–1649 (2011).

26. Uzair, M. et al. PERSISTENT TAPETAL CELL2 Is Required for Normal Tapetal Programmed Cell Death and Pollen Wall Patterning. Plant Physiol. 182, 962–976 (2020).

27. Li, N. et al. The rice tapetum degeneration retardation gene is required for tapetum degradation and anther development. Plant Cell 18, 2999–3014 (2006).

28. Kim, O. K., Jung, J. H. & Park, C. M. An Arabidopsis F-box protein regulates tapetum degeneration and pollen maturation during anther development. Planta 232, 353–366 (2010).

29. Béais, T. P. & Goldberg, R. B. A novel cell ablation strategy blocks tobacco anther dehiscence. Plant Cell 9, 1527–1545 (1997).

30. Vráblová, M., Vrábl, D., Sokolová, B., Marková, D. & Hronková, M. A modified method for enzymatic isolation of and subsequent wax extraction from Arabidopsis thaliana leaf cuticle. Plant Methods 16, (2020).

31. Vráblová, M., Vrábl, D., Hronková, M., Kubásek, J. & Šantrůček, J. Stomatal function, density and pattern, and CO2 assimilation in Arabidopsis thaliana tmm1 and sdd1-1 mutants. Plant Biol. 19, 689–701 (2017).

32. Ingouff, M. et al. Live-cell analysis of DNA methylation during sexual reproduction in Arabidopsis reveals context and sex-specific dynamics controlled by noncanonical RdDM. Genes Dev. 31, 72–83 (2017).

33. Cutler, S. R., Ehrhardt, D. W., Griffitts, J. S. & Somerville, C. R. Random GFP::cDNA fusions enable visualization of subcellular structures in cells of Arabidopsis at a high frequency. www.pnas.org. (2000).

34. Geldner, N. et al. Rapid, combinatorial analysis of membrane compartments in intact plants with a multicolor marker set. Plant J. 59, 169–178 (2009).

35. Fendrych, M. et al. Programmed cell death controlled by ANAC033/SOMBRERO determines root cap organ size in arabidopsis. Curr. Biol. 24, 931–940 (2014).

36. Wang, J. et al. A developmentally controlled cellular decompartmentalization process executes programmed cell death in the Arabidopsis root cap. Plant Cell 36, 941–962 (2024).

37. Olvera-Carrillo, Y. et al. A conserved core of programmed cell death indicator genes discriminates developmentally and environmentally induced programmed cell death in plants. Plant Physiol. 169, 2684–2699 (2015).

38. Van Durme, M. et al. Fertility loss in senescing Arabidopsis ovules is controlled by the maternal sporophyte via a NAC transcription factor triad. Proc. Natl. Acad. Sci. U. S. A. 120, (2023).

39. Gao, Z. et al. KIRA1 and ORESARA1 terminate flower receptivity by promoting cell death in the stigma of Arabidopsis. Nat. plants 4, 365–375 (2018).

40. Nelson, M. R. et al. A biomechanical model of anther opening reveals the roles of dehydration and secondary thickening. New Phytol. 196, 1030–1037 (2012).

41. Dobritzsch, S. et al. Dissection of jasmonate functions in tomato stamen development by transcriptome and metabolome analyses. BMC Biol. 13, 1–18 (2015).

42. Sanders, P. M., Bui, A. Q., Le, B. H. & Goldberg, R. B. Differentiation and degeneration of cells that play a major role in tobacco anther dehiscence. Sex. Plant Reprod. 17, 219–241 (2005).

43. Varnier, A. L., Mazeyrat-Gourbeyre, F., Sangwan, R. S. & Clément, C. Programmed cell death progressively models the development of anther sporophytic tissues from the tapetum and is triggered in pollen grains during maturation. J. Struct. Biol. 152, 118–128 (2005).

44. Papini, A., Mosti, S. & Brighigna, L. Programmed-cell-death events during tapetum development of angiosperms. Protoplasma 207, 213–221 (1999).

45. Dumais, J. & Forterre, Y. Vegetable dynamicks: The role of water in plant movements. Annu. Rev. Fluid Mech. 44, 453–478 (2011).

46. von Wangenheim, D. et al. Live tracking of moving samples in confocal microscopy for vertically grown roots. Elife 6, (2017).

47. R Core Team. R: A Language and Environment for Statistical Computing. (2023).

48. Postma, M. & Goedhart, J. Plotsofdata—a web app for visualizing data together with their summaries. PLoS Biol. 17, e3000202 (2019).

49. Wood, S. N. Fast stable restricted maximum likelihood and marginal likelihood estimation of semiparametric generalized linear models. J. R. Stat. Soc. Ser. B (Statistical Methodol. 73, 3–36 (2011).

50. Schindelin, J., et al. Fiji: An open-source platform for biological-image analysis. Nature Methods vol. 9 676–682 (2012).

51. Harrington, B. Inkscape.

