## Supplementary figures S1-15 for "Programmed cell death and stomatal density regulate anther opening in response to ambient humidity"

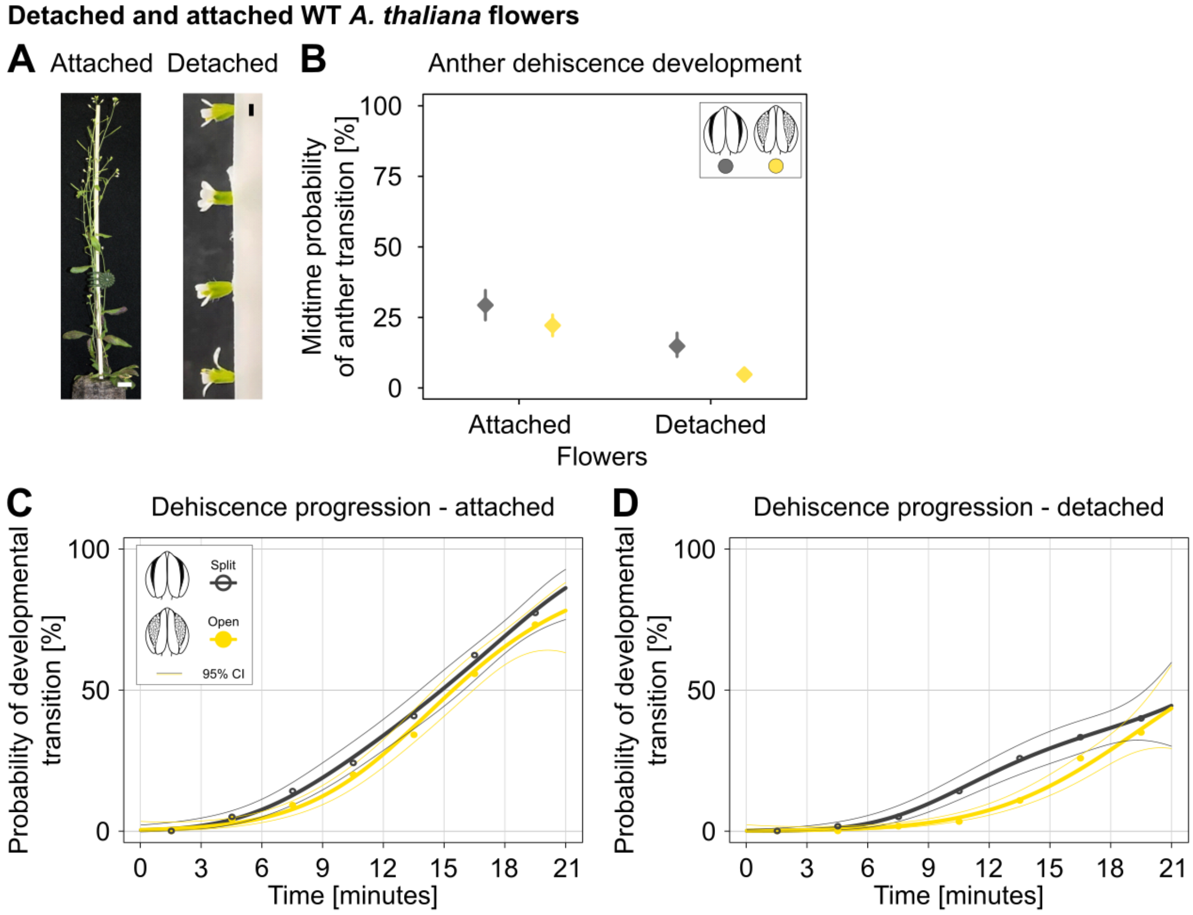


**Figure S1: Anther dehiscence is slower in *A. thaliana* flowers detached from plants. A.** Left: Still attached flowers, scale bar is 1 cm. Right: Detached flowers, scale bar is 5 mm. **B-D.** The development of 120 anthers from both attached and detached flowers was monitored for 21 minutes in 3-minute steps after short high humidity treatment was completed. Flower detachment significantly reduced both anther splitting rate (partially open anther in dark grey) and opening rate (fully open anther in yellow; GLM, $P < 2.2e\text{-}16^{***}$, 95% CI are shown). **B.** Rates from half-time of the measured period (10.5 minutes) are shown. **C**. Dehiscence progression of attached anthers. **D.** Anther dehiscence progression of detached flowers. **Legend:** Coloured points in C. and D. show measured proportions of respective developmental stages, logistic regression (GLM) is indicated by thick lines with 95% CI – thin lines (legend shown in figure C).

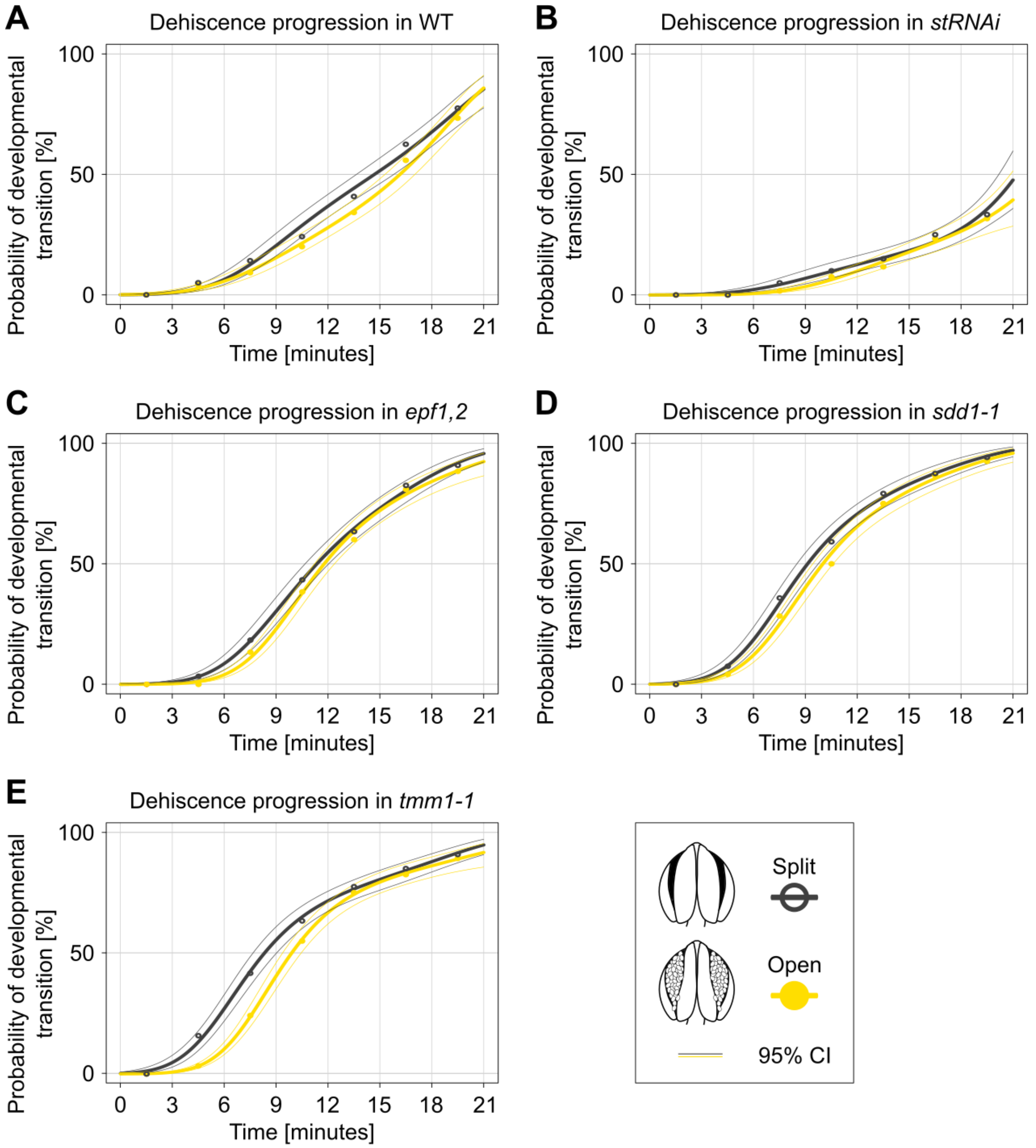


**Figure S2: Anthers of mutants with higher stomata density, *epf1,2*; *sdd1-1* and *tmm1-1*, split and open significantly faster compared to WT and *stRNAi*, a line with reduced stomata number. The probabilities of splitting (dark grey) and opening (yellow) are displayed, showing the dehiscence process over a 21-minute timeframe in 3-minute steps. A.** WT. **B.** *stRNAi*. Anther splitting and opening is notably decelerated. **C-E.** Anther splitting and opening are accelerated. **C.** *epf1,2.* **D.** *sdd1-1.* **E.** *tmm1-1*. **Legend:** The x-axis shows the time [minutes] and the y-axis indicates the probability rate of either splitting or opening. These graphs correspond with the bar plots from Figure 2. Coloured points show measured proportions of respective developmental stages, logistic regression (GLM) is indicated by thick lines with 95% CI – thin lines.

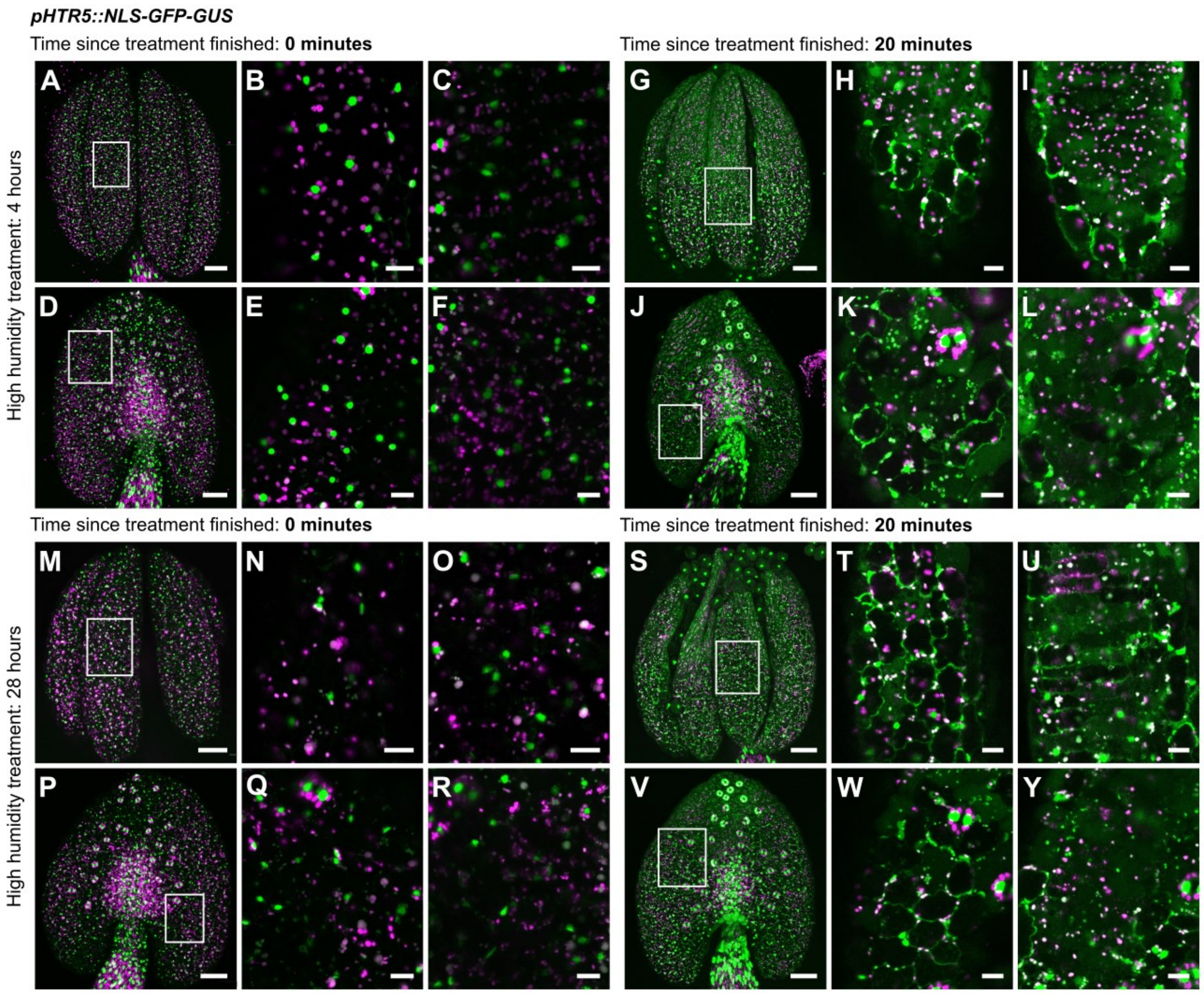


**Figure S3:** ***A. thaliana pHTR5::NLS-GFP-GUS*** **anthers stay closed when placed in high humidity conditions. Subsequent opening, when the treatment ends, is accompanied by programmed cell death (PCD).** The PCD progresses from the apex to the base, stomata also undergo PCD in the same direction. Nuclei are shown in green, autofluorescence in magenta. Once open anthers partially regain their original shape when placed in a liquid medium. **A-F.** Closed short-treated anthers. Both epidermis and endothecium are intact. **G-L.** Open anthers after a short treatment. Nuclei collapse in both, the epidermis and endothecium. **M-R.** Closed long-treated anthers. Most of the cells are still intact. **S-Y.** Open anthers after a long treatment, PCD progresses in the direction towards the base, and both epidermis and endothecium are affected. **Legend:** Flowers were exposed to high humidity for 4 or 28 hours and imaged 0 or 20 minutes after transfer to ambient humidity. For each group of figures, the upper row represents the adaxial side, and the lower row represents the abaxial side. From left to right: overall figures, epidermis details, endothecium details. The figures were obtained using Leica TCS SP8. Scale bars are 50 µm in overall view figures and 10 µm in close-up figures.

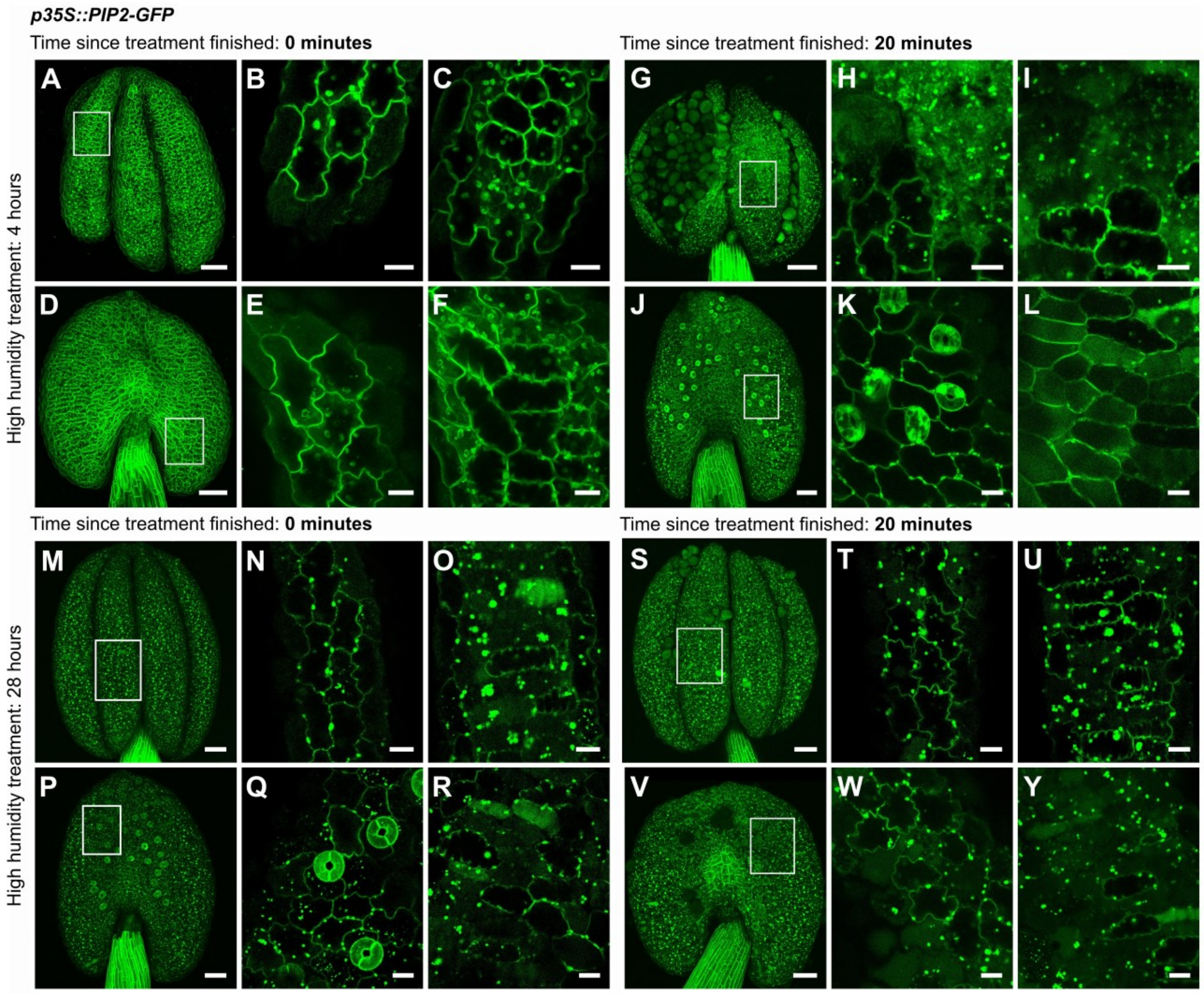


**Figure S4: Plasma membrane endodomain shedding is visible 20 minutes into anther opening.** *A. thaliana p35S::PIP2-GFP* anthers remain closed until the opening initiation, induced by short and long high humidity treatment. After 20 minutes, changes related to programmed cell death (PCD) occur on the plasma membrane (PM, shown in green). While open anthers can partly regain their shape when embedded in 2,5% low melting agarose, the PM changes persist. **A-F.** Closed anthers after a short treatment, PM in both epidermis and endothecium remains intact. **G-L.** Open anthers after a short treatment, PM in both epidermis and endothecium is already degraded. **M-R.** Closed anthers after a long treatment, PM is intact in the epidermis but not in the endothecium. **S-Y.** Open anthers after a long treatment, PM in both epidermis and endothecium is visibly affected. **Legend:** Flowers were exposed to high humidity for 4 or 28 hours and imaged 0 or 20 minutes after transfer to ambient humidity. For each group of figures: the upper row represents the adaxial side, and the lower row represents the abaxial side. From left to right: overall figures, epidermis details, endothecium details. The figures were obtained using Leica TCS SP8. Scale bars are 50 µm in overall view figures and 10 µm in close-up figures.

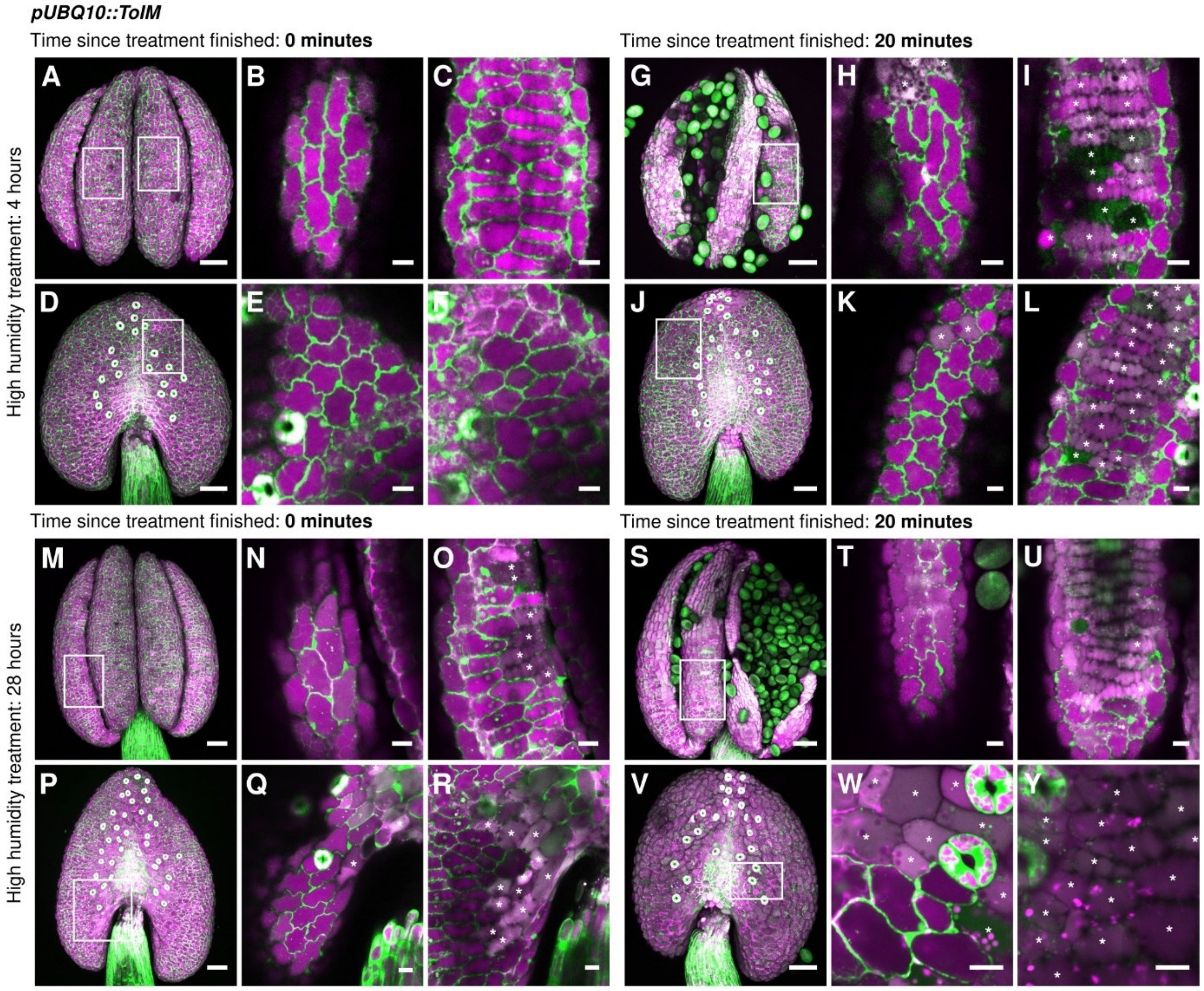


**Figure S5: The Tonoplast Integrity Marker (*A. thaliana pUBQ10::ToIM*) reveals progressing vacuole degradation within 20 minutes of anther opening.** The vacuole is represented by magenta and the cytoplasm by green. Upon vacuole bursting, the signals merge, resulting in a light pink to white signal – such cells are marked with asterisks. Anthers remain closed until the completion of both short and long high humidity treatments. However, in closed long-treated anthers at the time 0, vacuoles undergo degradation in the endothecium. Next, the progression of vacuole degradation and signal merging is observed in open anthers of both treated variants. **A-F.** Short-treated anthers with intact epidermis and endothecium. **G-L.** Open anthers after a short treatment, showing visible vacuole degradation predominantly in the endothecium. **M-R.** Closed anthers after a long treatment, with vacuole degradation already occurring in the endothecium. **S-Y.** Open anthers after a long treatment, displaying vacuole degradation in both epidermis and endothecium. **Legend:** Flowers were exposed to high humidity for 4 or 28 hours and imaged 0 or 20 minutes after transfer to ambient humidity. For each group of figures, the upper row represents the adaxial side, and the lower row the abaxial side. From left to right: overall figures, epidermis details, endothecium details. The figures were obtained using Leica TCS SP8. Scale bars are 50 µm in overall view figures, 10 µm in close-up figures.

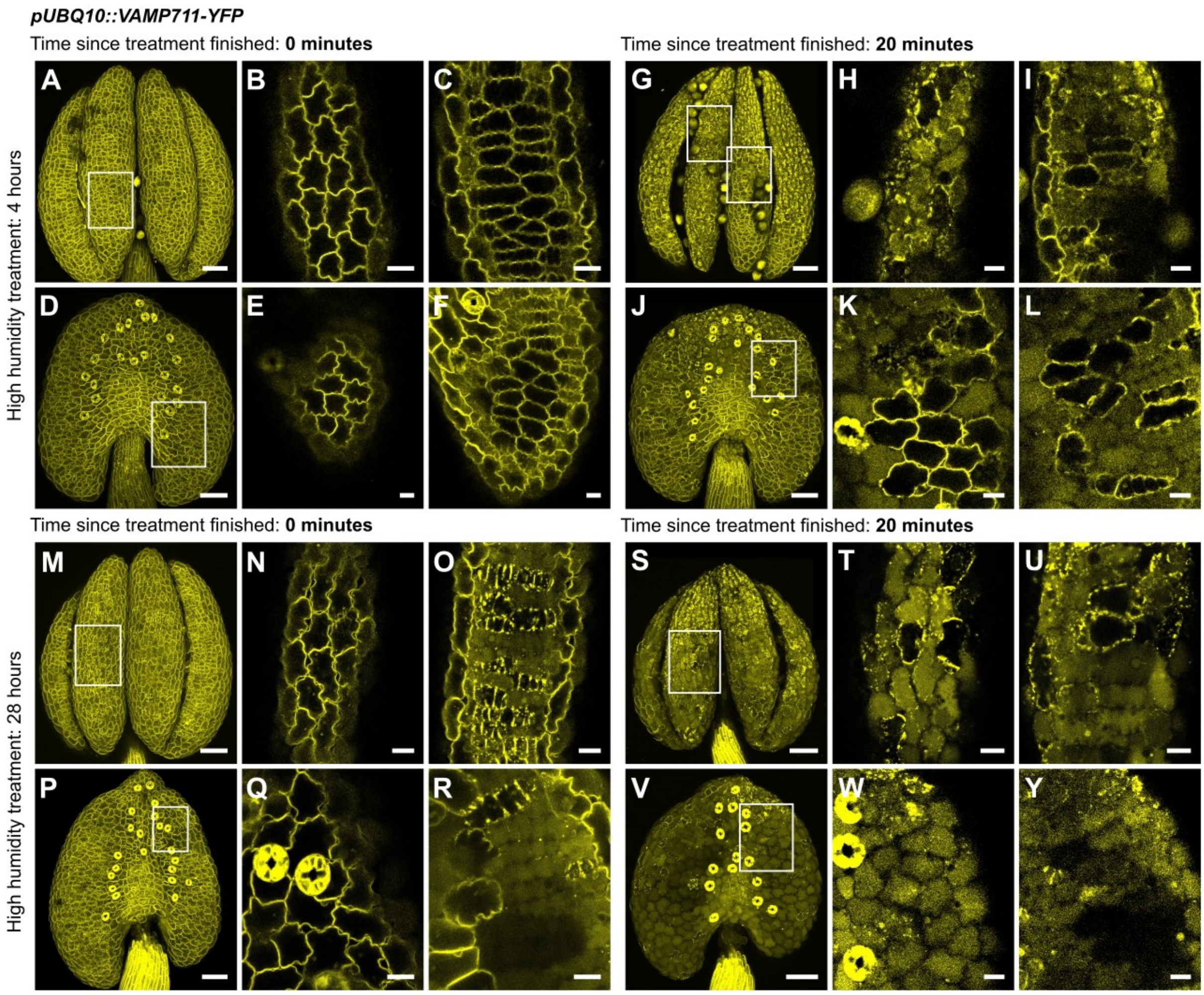


**Figure S6: Tonoplast marker shows endodomain shedding as anther opens.** *A. thaliana pUBQ10::VAMP711-YFP* anthers are closed until the opening starts when short and long high humidity treatment is finished. Tonoplast degradation is visible 20 minutes into anther opening (tonoplast shown in yellow). Open anthers can regain their shape when placed in 2,5% low melting agarose. **A-F.** Closed anthers after a short treatment; tonoplast in both epidermis and endothecium is intact. **G-L.** Open anthers after a short treatment; tonoplast in both epidermis and endothecium is already degraded, but tonoplast of some cells remains intact. **M-R.** Closed anthers after long treatment; tonoplast is intact in the epidermis but not in the endothecium. **S-Y.** Open anthers after a long treatment; tonoplast in both epidermis and endothecium is severely affected. **Legend:** Flowers were exposed to high humidity for 4 or 28 hours and imaged 0 or 20 minutes after transfer to ambient humidity. For each group of figures: the upper row represents the adaxial side, and the lower row represents the abaxial side. From left to right: overall figures, epidermis details, endothecium details. The figures were obtained using Leica TCS SP8. Scale bars are 50 µm in overall view figures, 10 µm in close-up figures.

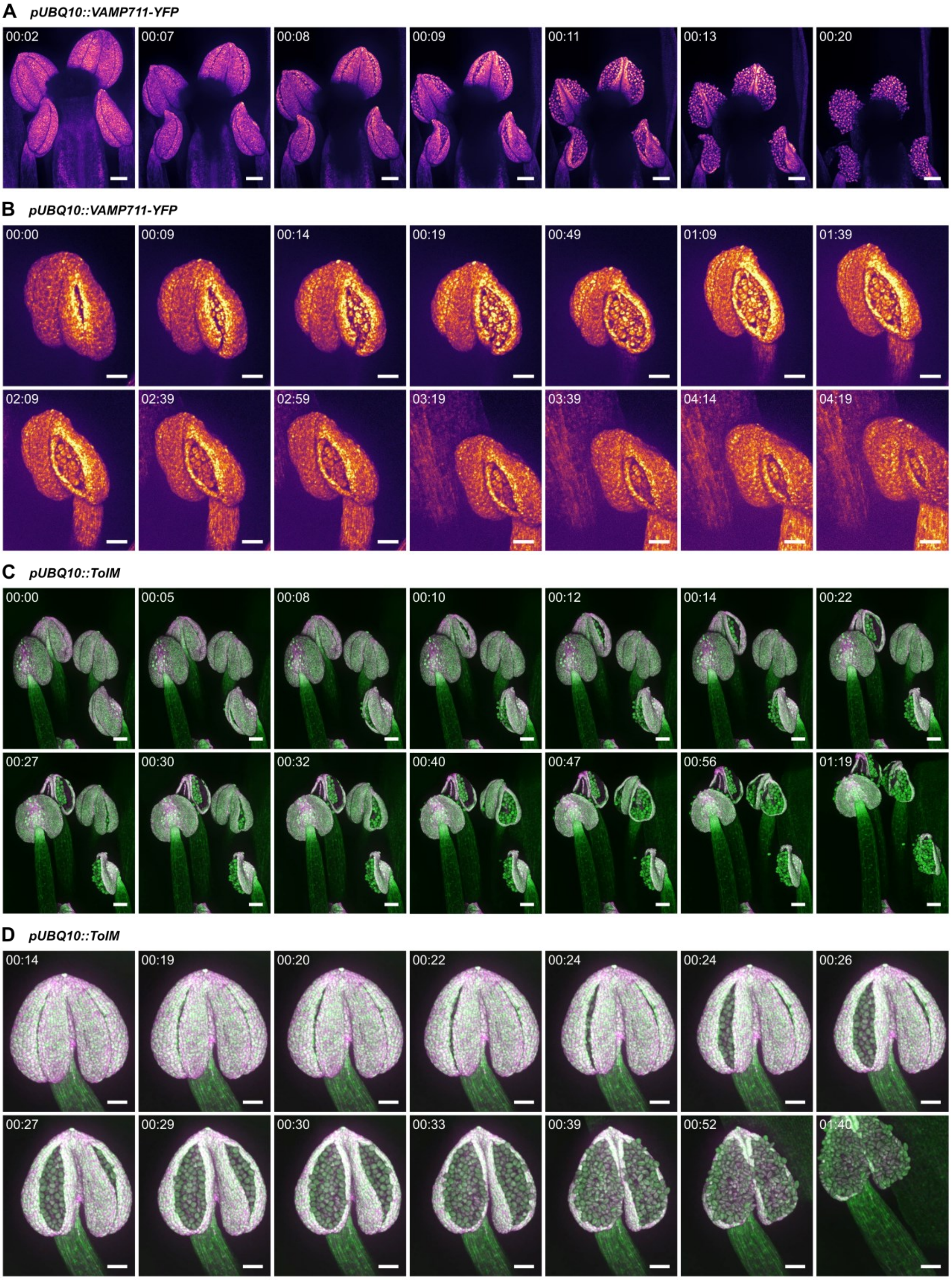


**Figure S7: Anther dehiscence progression can be captured using a spinning disc microscope with a dry mounting approach. This sample preparation enables a completely intact observation as the stamens remain attached to the plants the whole time. A-B.** *pUBQ10::VAMP711-YFP* line where tonoplast is shown in magenta-fire. **A.** Anther opening 20 minutes time lapse during which all anthers fully open. **B.** Anther dehiscence time-lapse, 4 hours and 19 minutes, reveals an opening initiation followed by anther closing and regaining the original shape. **C-D.** *pUBQ10::ToIM* line where vacuole is shown in magenta and cytoplasm in green. Signal merging, after the vacuole rupture, is visualised in light pink to white. **C.** The opening time-lapse, 1 hour and 19 minutes, shows the successive opening of all anthers. The filaments continue to elongate even after dehiscence completion. **D.** Time-lapse, 1 hour and 40 minutes, shows anther opening with subsequent filament elongation. All time-lapse series were obtained with a vertical stage (von Wangenheim *et al.*, 2017[46]) spinning disc microscope Zeiss Axio Observer.7/Yokogawa CSU-W1-T2 with a VS-HOM1000 excitation light homogenizer. The resolution was limited due to the dry mounting. Scale bars are 100 µm in A and C and 50 µm in B and D.

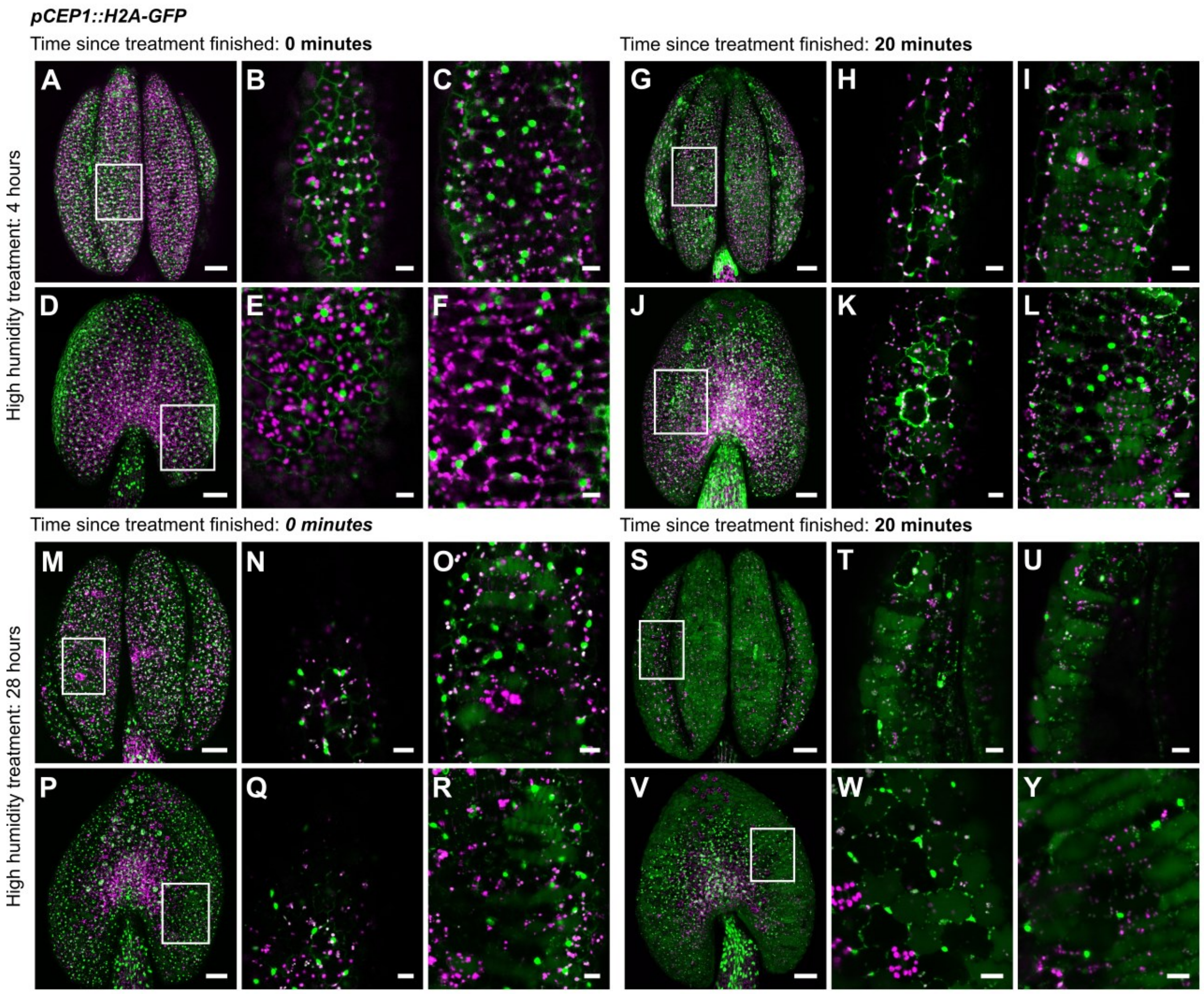


**Figure S8:** ***A. thaliana pCEP1::H2A-GFP*** **anthers show cells preparing for programmed cell death (PCD), with expression visible in epidermis and endothecium.** Nuclei are in green, autofluorescence in magenta. Anthers remain closed during short and long high humidity treatments, even when the endothecium undergoes PCD during the long treatment. Following treatment completion, anthers open, and cells undergo PCD as the signal spread to the whole cells volume. Already open anthers can partly regain their shape when embedded in 2,5% low melting agarose. **A-F.** Closed anthers after a short treatment; both epidermis and endothecium are fully viable. **G-L.** Open anthers after a short treatment; cells begin undergoing PCD, with some remaining intact. **M-R.** Closed anthers after a long treatment; epidermal cells are mostly intact, while PCD in the endothecium has already commenced. **S-Y.** Open anthers after a long treatment; both epidermis and endothecium exhibit signs of PCD. **Legend:** Flowers were exposed to high humidity for 4 or 28 hours and imaged 0 or 20 minutes after transfer to ambient humidity. For each group of figures, the upper row represents the adaxial side, and the lower row represents the abaxial side. From left to right: overall figures, epidermis details, endothecium details. The figures were obtained using Leica TCS SP8. Scale bars are 50 µm in overall view figures, 10 µm in close-up figures.

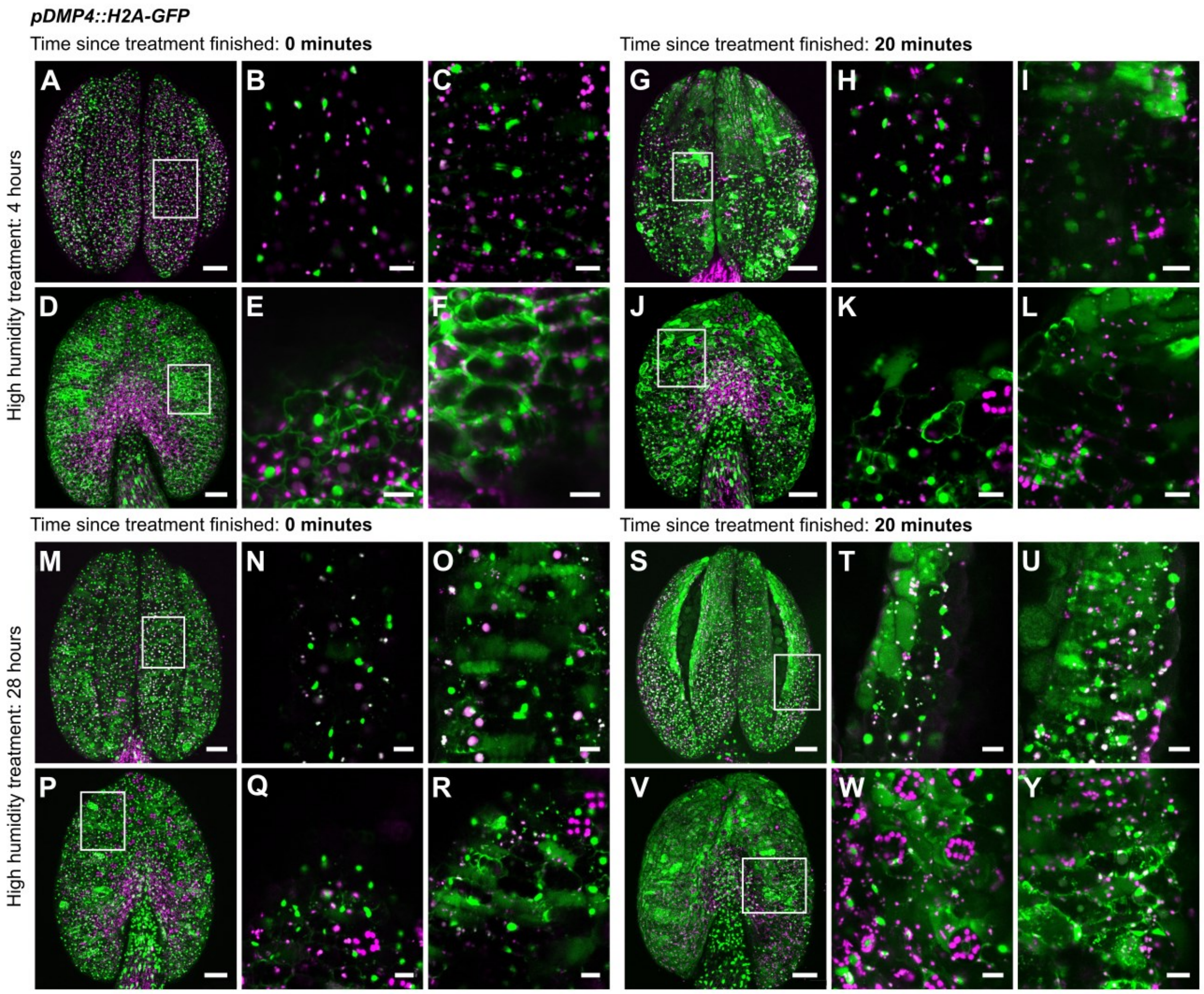


**Figure S9:** ***A. thaliana pDMP4::H2A-GFP*** **anthers cells prepare for programmed cell death (PCD) before opening and then undergo PCD as the anthers open.** High humidity treatments prevent anthers from opening. GFP is shown in green and is restricted primarily to the nuclei in intact cells. Autofluorescence is shown in magenta. As anthers open, the PCD progresses and the nuclei collapse, the signal diffuses and fills the cells. Once open, anthers can regain their shape when placed in 2,5% low melting agarose, but the changes on the cell level are permanent. **A-F.** Closed short-treated anthers; cells are still intact. **G-L.** Open anthers after a short treatment; PCD progresses in both epidermis and endothecium, the apex possesses more dead cells than the base. **M-R.** Closed long-treated anthers; cells are intact in the epidermis; the endothecium contains already dead cells. **S-Y.** Open anthers after a long treatment; most of the cells are dead except for the connective on the abaxial side. PCD again proceeds from the anther apex to the base. **Legend:** Flowers were exposed to high humidity for 4 or 28 hours and imaged 0 or 20 minutes after transfer to ambient humidity. For each group of figures: the upper row represents the adaxial side, and the lower row represents the abaxial side. From left to right: overall figures, epidermis details, endothecium details. The figures were obtained using Leica TCS SP8. Scale bars are 50 µm in overall view figures, 10 µm in close-up figures.

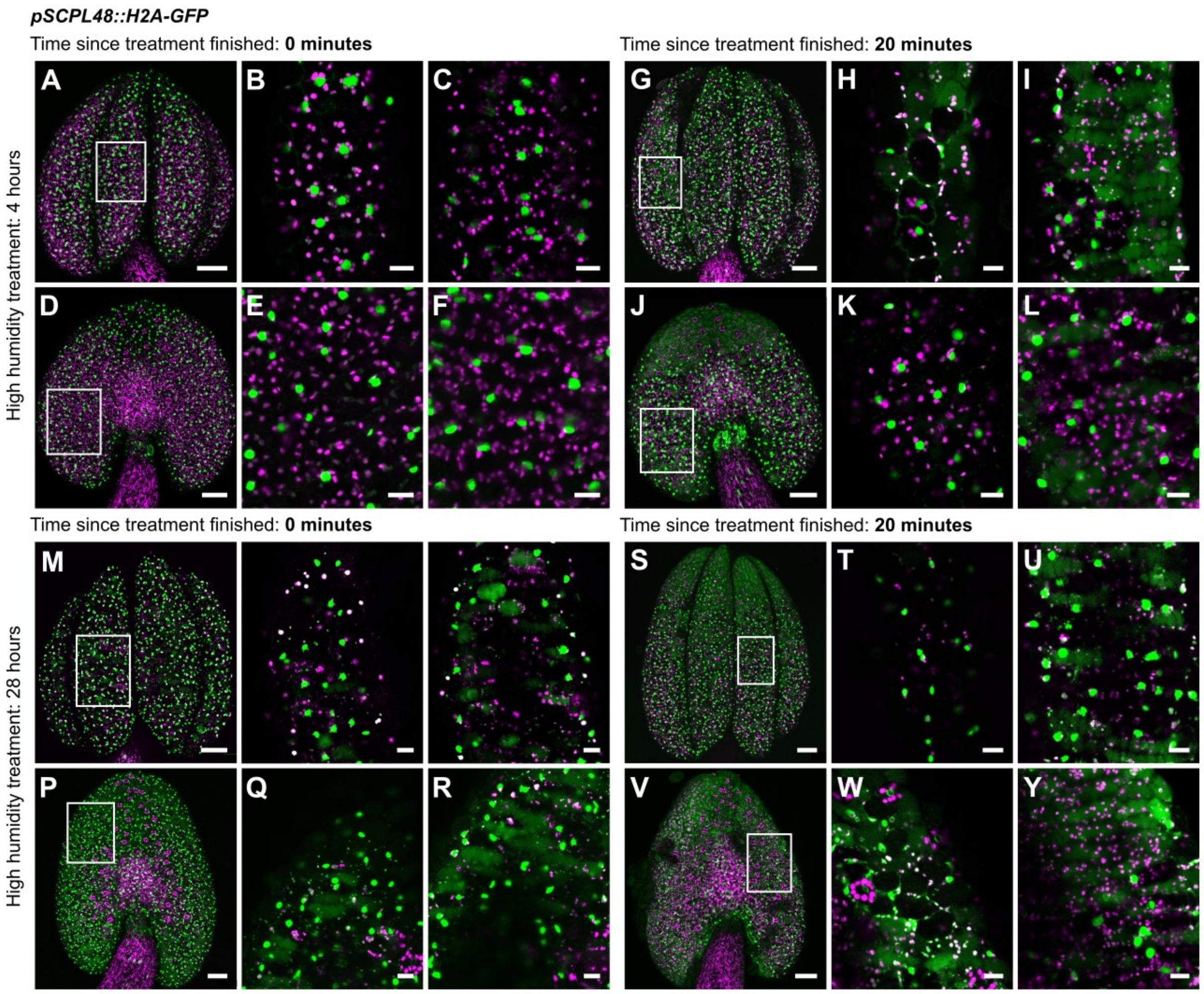


**Figure S10: Anthers of *A. thaliana pSCPL48::H2A-GFP* are pre-programmed for cell death (PCD), which is facilitated only once high humidity treatments end and anthers open.** GFP, shown in green, is localised to the nuclei of living cells of closed anthers. During the anther opening, nuclei collapse, cells die, and the signal fills the cells. Autofluorescence is in magenta. Already open anthers can regain their original shape when placed in 2,5% low melting agarose, but the changes on the cell level are definite. **A-F.** Closed short-treated anthers; cells are viable. **G-L.** Open anthers after a short treatment; PCD progresses from the apex to the base, and both epidermis and endothecium are affected. **M-R.** Closed long-treated anthers; epidermal cells are intact, but some endothecial cells are already dead. **S-Y.** Open anthers after a long treatment; cells die in the direction from the apex to the base. Connective tissue stays viable. **Legend:** Flowers were exposed to high humidity for 4 or 28 hours and imaged 0 or 20 minutes after transfer to ambient humidity. For each group of figures: the upper row represents the adaxial side, and the lower row represents the abaxial side. From left to right: overall figures, epidermis details, endothecium details. The figures were obtained using Leica TCS SP8. Scale bars are 50 µm in overall view figures, 10 µm in close-up figures.

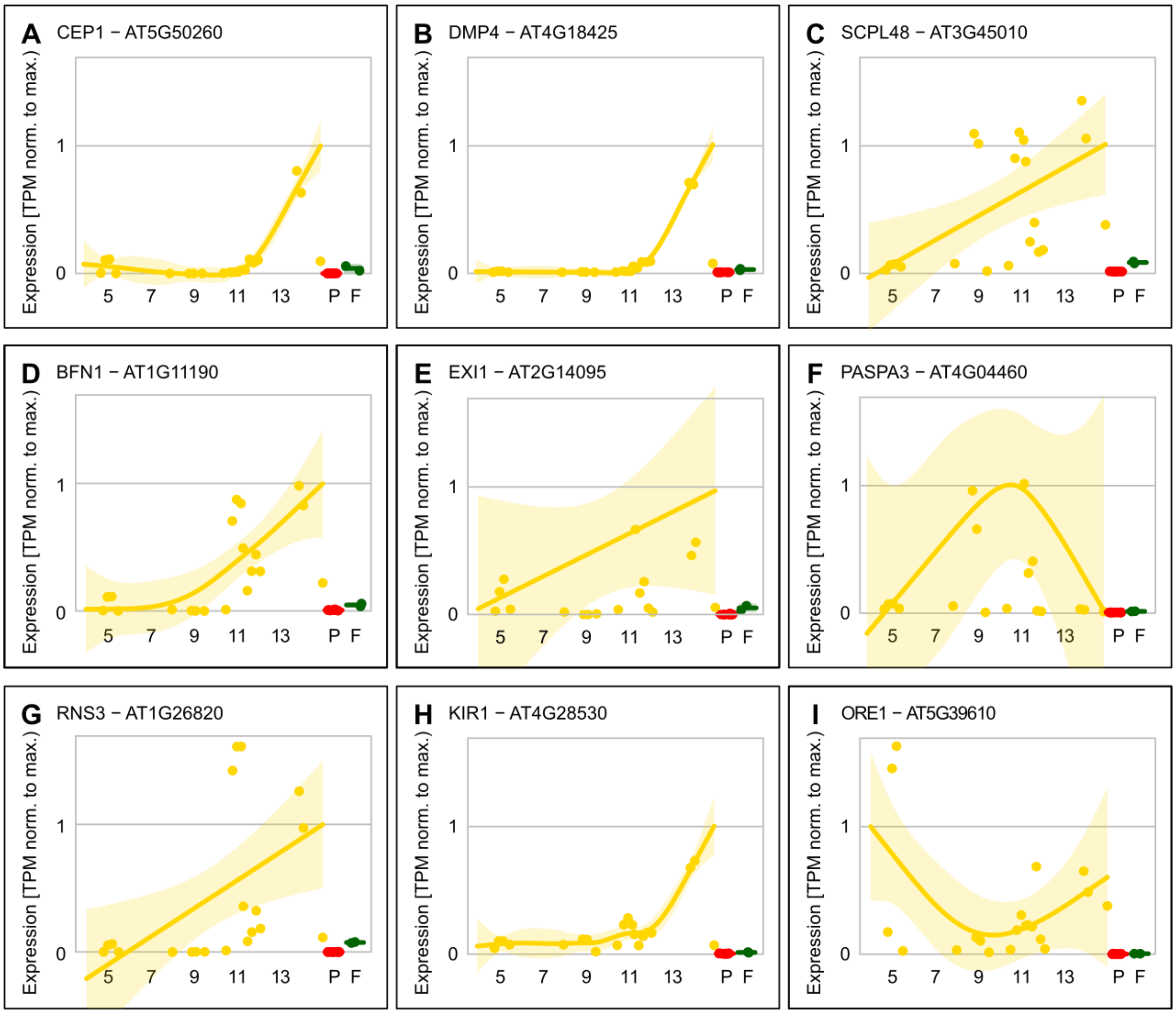


**Figure S11: The expression of studied PCD-related genes *BFN1*, *CEP1*, *DMP4*, *EXI1*, *KIR1*, *ORE1*, *RNS3* and *SCPL48* increases in later anther development. On the contrary, PASPA3 expression decreases.** Publicly available RNA-seq data of 12 different developmental phases of anther, 3 independent transcriptomics of mature pollen and one transcriptomic of filament (Table S3-4), were quantified using Kallisto 0.48.0 against Araport11 representative CDS model and TPM (transcripts per million) were fitted by GAM (for anther stages) or used for median computation (pollen, filament). In each graph, individual gene transcription (as relative value to fitted maximum) with 95% confidence intervals is shown. Numbers on the x-axis stand for anther development stage according to Sanders *et al.*, 1999[8], P for mature pollen and F for filaments.

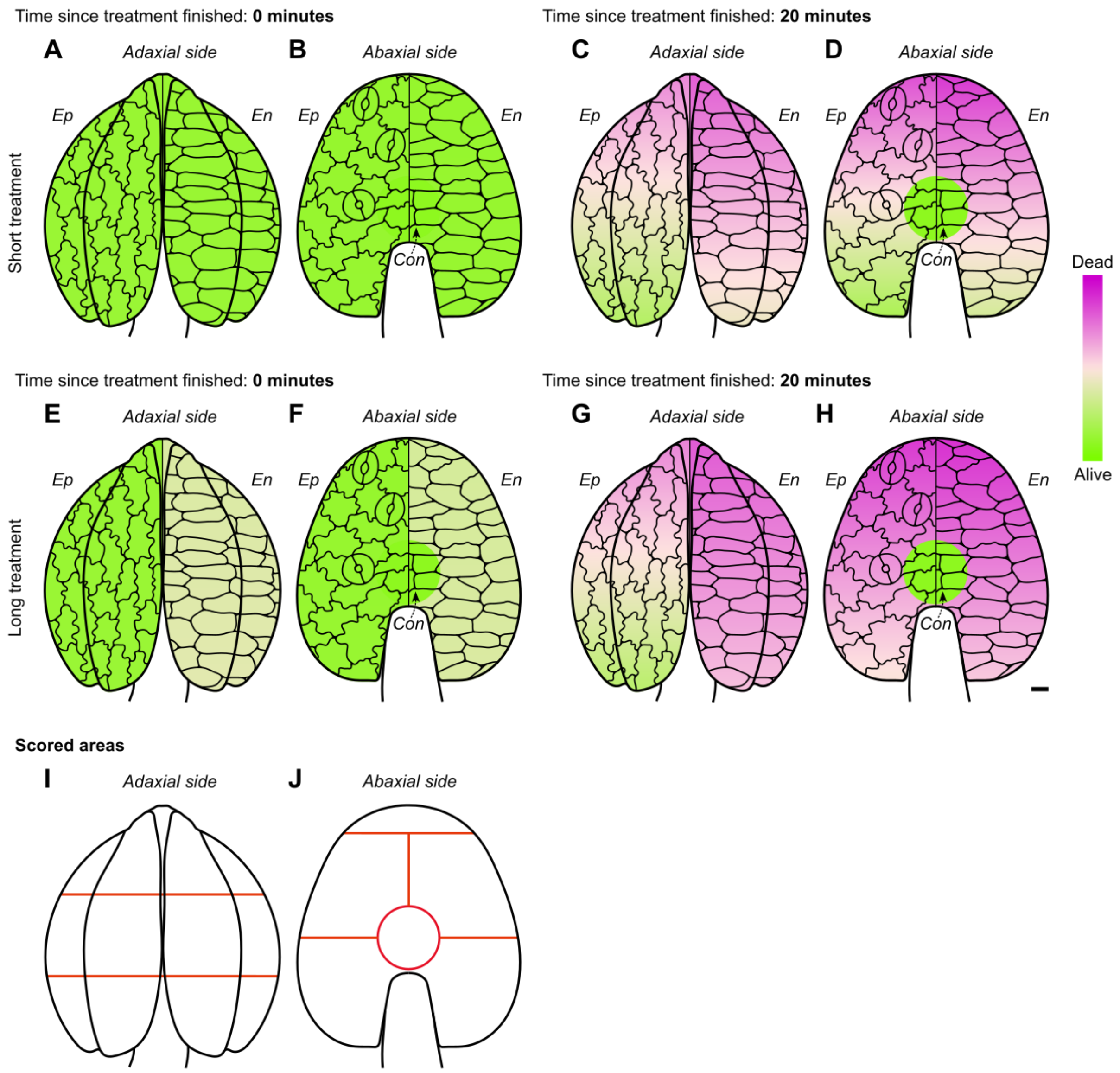


**Figure S12: Cell death progresses with time in the direction from the anther apex to the base.** In total, 112 anthers were analysed from following fluorescent marker lines: *pHTR5::NLS-GFP-GUS*, *p35S::PIP2-GFP, pUBQ10::ToIM*, *pUBQ10::VAMP711-YFP*, *pCEP1::H2A-GFP*, *pDMP4::H2A-GFP* and *pSCPL48::H2A-GFP*. Anthers were exposed to short (4 hours) or long (28 hours) high humidity treatment and imaged 0 or 20 minutes after the treatments finished. **A-B.** Short-treated anthers, adaxial (AD) and abaxial (AB) sides, at the time 0. Both epidermis and endothecium are fully viable. **C-D.** Short-treated anthers, AD and AB sides, 20 minutes after the treatment. Epidermal cells stay intact longer than endothecial cells on both sides. Anther apex contains more dead cells than the base. Connective cells are still viable. **E-F.** Long-treated anthers, AD and AB sides, at the time 0. The epidermis is intact, but endothecium undergoes PCD except for connective tissue. **G-H.** Long-treated anthers, AD and AB sides, 20 minutes after the treatment. The PCD has a similar pattern to the short-treated anthers but progresses faster. Connective cells are viable. **I-J.** Anthers were divided into several regions indicated by the red lines: apex, middle and base. Also, the connective area was distinguished at the abaxial side. These areas were scored according to the dead cell content, > 10 %, 10-40 %, 40-60 %, 60-90 % and > 90 %. Red lines indicate areas that we scored. **Legend:** The data were analysed using beta regression in R (package **betareg**). Magenta represents the completely dead cells, lawn green the fully living cells and mistyrose represents the transition between dead and living tissue. Each anther scheme has an epidermis (*Ep*) on the left side and an endothecium (*En*) on the right side. The cell schemes represent the assigned tissue of respected anther sides, AD and AB. *Con.* indicates connective tissue. The size of the cells in comparison to the anther scheme size is approximately 4 times bigger. The scale bar is 10 μm.

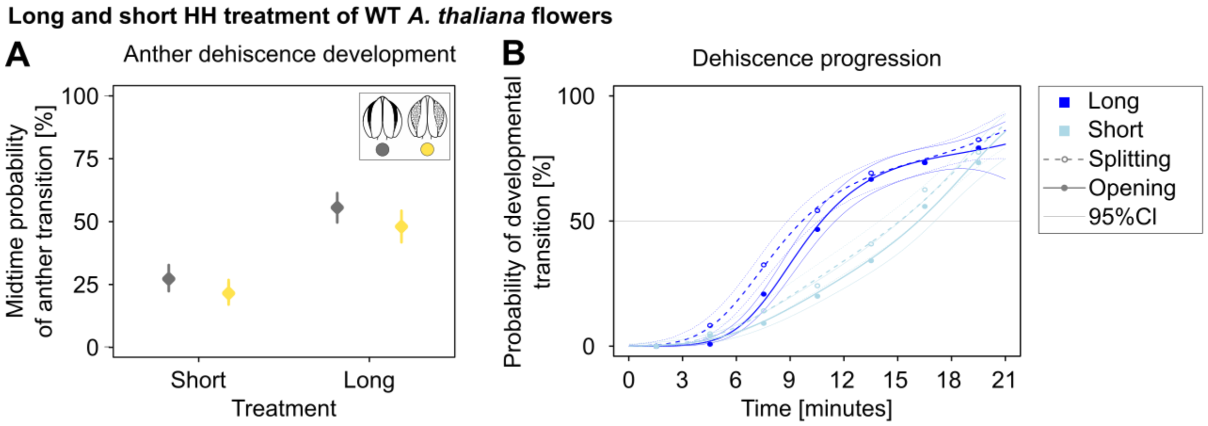


**Figure S13: Anther dehiscence is faster in *A. thaliana* anthers exposed to 24 hours of high humidity treatment (long HH treatment) compared to short-treated anthers (4 hours). A.** Long treatment significantly increases both anther splitting (dark grey) and opening (yellow; GLM, $P < 2.2e\text{-}16^{***}$, 95% CI are displayed). Rates from half-time of the measured period (10.5 minutes) are shown. **B.** The curves show splitting and opening occurring earlier in long treated anthers. Splitting (dashed line) and full opening (solid line) in anthers exposed to long HH treatment (dark blue) compared to the short treatment (light blue). 95% CI are shown.

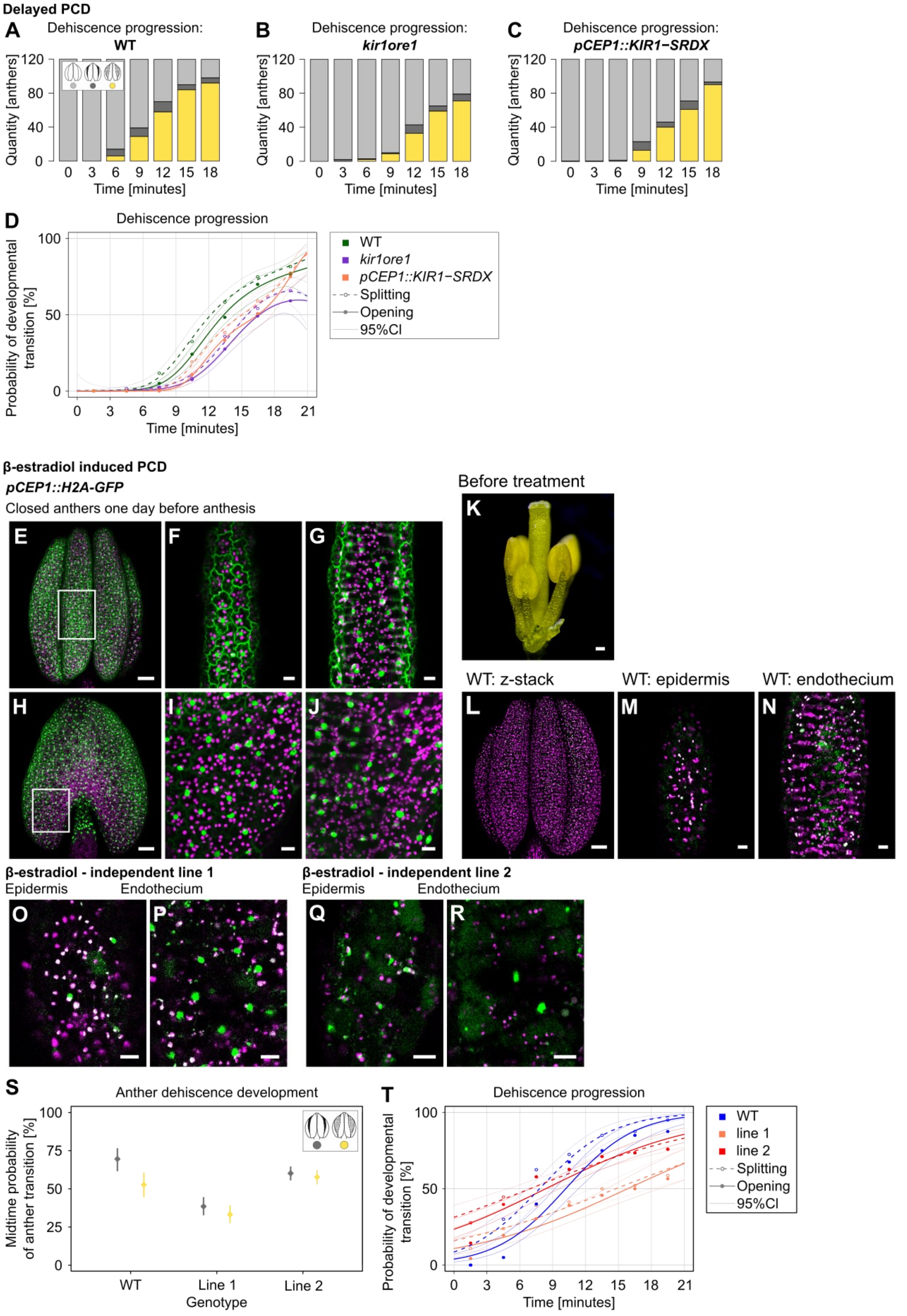


**Figure S14: Genetic manipulation of PCD changes the anther splitting and opening timing. A-D.** ***pCEP1::KIR1-SRDX*** **and** ***kir1ore1*** **anthers splitting and opening is decelerated when compared to**

**wild-type (WT).** Anther dehiscence was measured over 21 minutes in 3-minute steps, all graphs show the dehiscence progression. **A.** WT anther dehiscence, in the midtime, 39 out of 120 are split and open. **B.** *kir1ore1* dehiscence progression, 10 out of 120 are split and open in the midtime. **C.** *pCEP1::KIR1-SRDX* anther dehiscence progression, 23 out of 120 split and open in the midtime. **D.** Probability of anther splitting and opening shows reduced anther dehiscence progression in *pCEP1::KIR1-SRDX* and *kir1ore1* compared to WT (GLM, $P$ < 2.2e-16***, rates and their 95% CI are shown). Legend in the graph. **E-T. Opening despite high humidity treatment occurs in *pCEP1::XVE≫KIR1-GFP* anthers after PCD is stimulated. E-J.** Expression in the whole tissue occurs in the *pCEP1::H2A-GFP* anthers 24 hours before dehiscence finalisation. This serves as a control that CEP1 is expressed at the time of β-estradiol treatment. **E.** Adaxial side z-stack. **F.** Epidermis on the adaxial side. **G.** Endothecium on adaxial side. **H.** Abaxial side z-stack. **I.** Epidermis on the abaxial side. **J.** Endothecium on the abaxial side. **K.** *pCEP1::XVE≫KIR1-GFP* flower at the beginning of β-estradiol treatment, 24 hours before anthesis. To ensure the β-estradiol solution accessibility, the pistil, sepals and petals are removed (the pistil is kept only for the picture to compare the unmatured stamens length to the pistil length). **L-M.** WT with no induced expression after β-estradiol treatment. **L.** Adaxial side of the anther, z-stack. **M.** Detail of epidermis. Round bodies with overlapping emission spectra (green and magenta resulting in light pink to white) are oil bodies. They are also visible in bright-field z-stack as they are dense and move rapidly in contrast to nuclei. **N.** Detail of endothecium. **O-P.** *pCEP1::XVE≫KIR1-GFP* (independent line 1) 24 hours after β-estradiol induction. **O.** Expression in epidermal cells. **P.** Expression in endothecial cells. **Q-R.** *pCEP1::XVE≫KIR1-GFP* (independent line 2) 24 hours after β-estradiol induction. Tissue already undergoes PCD. **Q.** Expression in epidermal cells. **R.** Expression in endothecial cells. **S-T.** Dehiscence progression was measured in inducible lines and WT as a control over 21 minutes in 3-minute steps (GLM, $P$ < 2.2e-16***, rates and their 95% CI are shown). **S.** Midtime anther splitting and full opening probability differed among all the lines. Legend in the figure. **T.** Probability of anther splitting and full opening shows that anthers of inducible lines open despite the high humidity. However, the splitting and opening of WT anthers is more rapid over time course. **Legend:** All the fluorescent figures were obtained using Leica TCS SP8. The scale bar in z-stack figures (E, H, L) is 50 μm. The scale bar in close up-figures (F-G, I-J, M-N, O-R) is 10 μm. Figure K was captured with a Nikon D3200 camera and stereomicroscope STM 822 5410, with a scale bar of 100 μm.

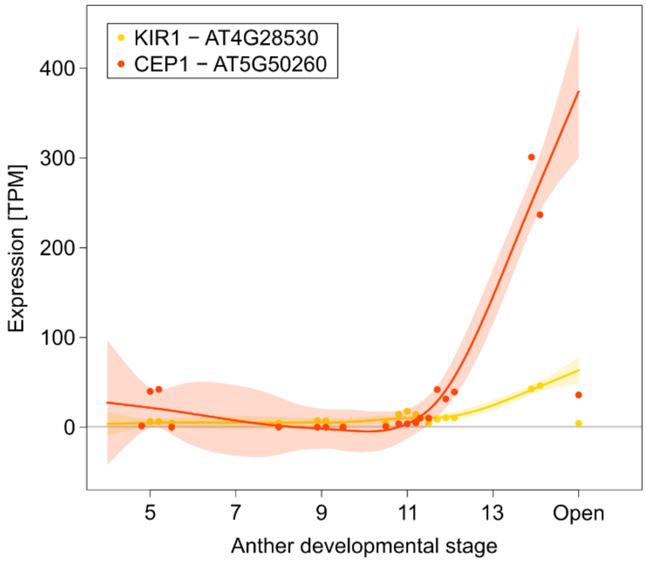


**Figure S15: The expression of PCD-related genes, *CEP1* and *KIR1* from the inducible overexpression line *pCEP1::XVE≫KIR1-GFP* (Gao *et al.*, 2018), increases prior to anther dehiscence.** Publicly available RNA-seq data of different developmental phases of anther (Table S3-4) were quantified using Kallisto 0.48.0 against Araport11 representative CDS model and TPM (transcripts per million, y-axis) were fitted by GAM. In the graph, individual genes transcription (expression curves were not normalised to the min-max range) with 95% CI are shown. Individual libraries were not normalized before quantification, quantile-normalized expression profiles gave almost identical expression pattern. Numbers on the x-axis stand for anther development stage according to Sanders *et al.*, 1999[8].
